# CNS Myelin Sheath Lengths Locally Scale to Axon Diameter via Piezo1

**DOI:** 10.1101/2025.11.18.689045

**Authors:** Amanda R. Young, Ryan W. Lewis, Martha Cash, Beckam Polis, Myah Zalusky, Jacob Reyngoudt, Avipsha Datta, Marie E. Bechler

**Affiliations:** Department of Cell and Developmental Biology, SUNY Upstate Medical University, Syracuse, NY; Department of Neuroscience and Physiology, SUNY Upstate Medical University, Syracuse, NY

**Keywords:** myelin sheaths, internode length, mechanosensing, axon caliber, axon diameter, intrinsic myelination, mechanosensitive ion channel, oligodendrocytes, Piezo1

## Abstract

Myelin sheath lengths vary by an order of magnitude in the central nervous system (CNS) and tune the timing of neuronal signaling. Thus, variation in myelin sheath length has been proposed to coordinate the timing of neuronal signaling to ultimately impact behavior. The mechanisms to establish myelin sheath length are unknown. For decades, reports have documented that in vivo myelin sheath size scales with the diameter of the ensheathed axon. We previously demonstrated diameter is sufficient to instruct myelin sheath lengths formed by oligodendrocytes using a synthetic axon culture system. The mechanisms of oligodendrocyte diameter-sensing and its translation into sheath elongation are still unknown. Here we demonstrate that diameter-sensing and sheath length is locally regulated: each individual myelin sheath responds to the underlying fiber diameter. We uncover a novel mechanism for scaling myelin sheath length to fiber diameter, through mechanosensitive ion channel Piezo1. In vivo, Piezo1 impacts the elongation of myelin sheaths on large diameter axons, recapitulating our in vitro results. Yet, surprisingly, there is no impact on myelin thickness with conditional Piezo1 loss. We propose Piezo1 provides a mechanism to establish hard-wired myelin sheath patterns, where oligodendrocytes transduce axon diameter into generating myelin segments with vastly different lengths.

## Introduction

Myelin sheaths support the health and energy efficiency of axons^1^ as well as impact the speed of action potential propagation along axons. Signalling speed in myelinated axons greatly depends on the physical properties of the myelin sheaths, especially sheath length (i.e., internodal distance)^2,3^. Strikingly, the CNS displays an order of magnitude variation in myelin sheath lengths, from 10s to 100s of microns in length^4,5,6^. Together, this has spurred the hypothesis that variation in myelin sheath lengths is critical to CNS function by precisely adjusting neuronal signaling speeds to facilitate signal synchronization within neuronal circuits^7,8^. Indeed, tuning myelin sheath size to coordinate signal timing may be a fundamental mechanism to enable behavior^9,10,11^ and can be perturbed in an array of neurodevelopmental^12,13^ and neurodegenerative conditions^14^. What governs how CNS myelin sheath lengths are initially established is still unclear.

For decades, in vivo studies have demonstrated a nearly linear positive correlation between myelin sheath lengths and axon diameters for the majority of myelinated axons^15,16,17^. How oligodendrocytes scale myelin sheath lengths with axon diameter is unknown. Studies have pointed to a major role for diameter sensing in both onset of myelin sheath formation as well as myelin sheath lengths. Reductionist approaches with synthetic axons (fibers that mimic the size and shape of axons) demonstrated that a threshold fiber diameter is sufficient to trigger myelination^18^. Complementary in vivo studies also showed that increased axon diameter can stimulate unmyelinated axons to become myelinated^19^. Consistent with diameter being sufficient to stimulate myelination, in vivo data has demonstrated myelination can occur independently of active neuronal signals^20,21,22,23^. Direct evidence that diameter also instructs sheath length was demonstrated with isolated oligodendrocytes on synthetic axons, where myelin sheath lengths scaled with the diameter of synthetic axons^24^. Taken all together, these results led to the proposed model that oligodendrocytes sense and use axon diameter as a cue to initiate oligodendrocytes’ intrinsic ability to form myelin sheaths and establish CNS myelin sheath lengths^25^. In such a model, the molecular pathways sensing and responding to axon diameter are anticipated to be some of the main drivers of neuronal-activity-independent myelin sheath growth. In this work, we sought to identify how diameter (axon or microfiber) is sensed as a physical cue and translated into control of myelin sheath lengths.

Mechanotransduction, the ability to sense and convert physical cues into cellular responses, is an increasingly appreciated but understudied property of many cell types, including oligodendrocyte lineage cells. Physical cues such as extracellular stiffness affect oligodendrocyte lineage cell differentiation and can have opposing effects on myelination. Increased substrate stiffness promotes changes to transcription factor localization, can activate transcriptional and/or epigenetic changes, as well as alter oligodendrocyte lineage differentiation^26,27,28,29,30^. Multiple mechanotransduction pathways including LINC complex, YAP/TAZ, integrin, and mechanosensitive ion channel signalling have all been implicated in oligodendrocyte maturation^26,28,31,32,33,34^. Interestingly, mechanotransduction during myelin formation appears to be regulated opposingly to oligodendrocyte differentiation: pliable substrates promote myelin membrane wrapping of synthetic axons while wrapping is hindered by stiff substrates^30^. Emerging evidence of mechanotransduction in oligodendrocytes highlights the important, open question of how differentiated oligodendrocytes integrate physical cues to regulate myelin sheath formation and properties. The mechanotransduction pathways that drive the formation of spirally wrapped myelin membranes are still unknown.

Here we investigate how oligodendrocytes sense axon diameters and translate this physical signal into myelin sheath lengths. We establish that diameter sensing is transduced at the individual myelin sheath level and identify a mechanosensitive protein responsible for myelin sheath elongation on large diameter axons.

## Results

### Myelin Sheath Lengths Scale Locally with Axon Diameter

To determine the mechanisms that oligodendrocytes use to scale myelin sheath growth (lengths) with the physical cue of diameter, we have taken advantage of oligodendrocyte cultures with electrospun microfibers^35^. These microfibers mimic the size and shape of CNS axons and are sufficient to promote isolated rodent oligodendrocytes to form compact, multilamellar myelin sheaths that mirror myelin sheath lengths observed in vivo^24^. An important strength of this model is the ability to tightly control culture conditions and fiber diameter, thereby eliminating any influences from the CNS biochemical milieu. Oligodendrocytes cultured on uniform microfibers of specific diameter ranges (0.5-1 micron, 1-2 micron, or 2-4 micron) scale myelin sheath lengths to the diameter of the fibers^24^, mimicking observed in vivo correlations of myelin sheath length with axon diameter^15,16,17^. This neuron-free system is ideal for elucidating how axon diameter is translated into myelin sheath lengths, because it enables testing the contribution of diameter as the sole variable during the 3-D process of myelin sheath formation.

First, we addressed whether oligodendrocytes respond to fiber diameter on a whole-cell level or if every individual nascent myelin sheath locally regulates length in response to diameter. Mechanotransduction in oligodendrocyte lineage cells has previously been shown to involve YAP/TAZ or LINC complex signalling that mediates transcriptional or epigenetic changes^26,30,31^. In contrast to this, individual oligodendrocytes can generate myelin sheaths that vary in size, suggesting local regulation^4,36,37^. To distinguish between global (e.g., transcriptional) or local responses, we modified the design of our microfiber system to generate mixed diameter microfiber scaffolds (Figure 1 A, B). Mixed diameter microfibers enabled assessment of myelin sheaths formed by single oligodendrocytes simultaneously in contact with both small and large diameter fibers. We predicted that if signalling led to whole-cell responses, the myelin sheaths lengths would no longer scale with diameter in mixed diameter fiber cultures. However, if sheath length is locally regulated, we predicted that sheath lengths would scale with diameter, equivalent to myelin sheaths in uniform-diameter microfiber cultures (Figure 1 A). Myelin sheath length in response to underlying fiber diameter was assessed from primary rat oligodendrocytes cultured on mixed diameter microfibers for 14 days, when maximum sheath lengths are formed^24^. Each myelin sheath length and the corresponding fiber diameter was measured from single oligodendrocytes simultaneously in contact with both large and small diameter fibers, using confocal micrographs (Figure 1 B, Figure S1). We observed that myelin sheath lengths increased with the underlying fiber diameter (Figure 1 C), supporting that each nascent myelin sheath length locally scales with the diameter of the fiber. To confirm whether myelin sheath lengths are locally controlled in each nascent myelin sheath on a larger scale, we binned a total of 127 myelin sheath lengths formed on small diameter (0.5-2.5 microns) and large diameter fibers (2.6-6 microns, Figure S1 A,B). We compared the binned myelin sheath length distribution from mixed diameter cultures against the distribution of myelin sheath lengths in uniform-diameter microfibers. The distribution of myelin sheath lengths between mixed diameter and uniform diameter microfiber cultures were statistically equivalent (Figure 1 D, data shown per replicate in Figure S1 C), consistent with a model where myelin sheath lengths are controlled at the level of individual sheaths.

**Figure1.**
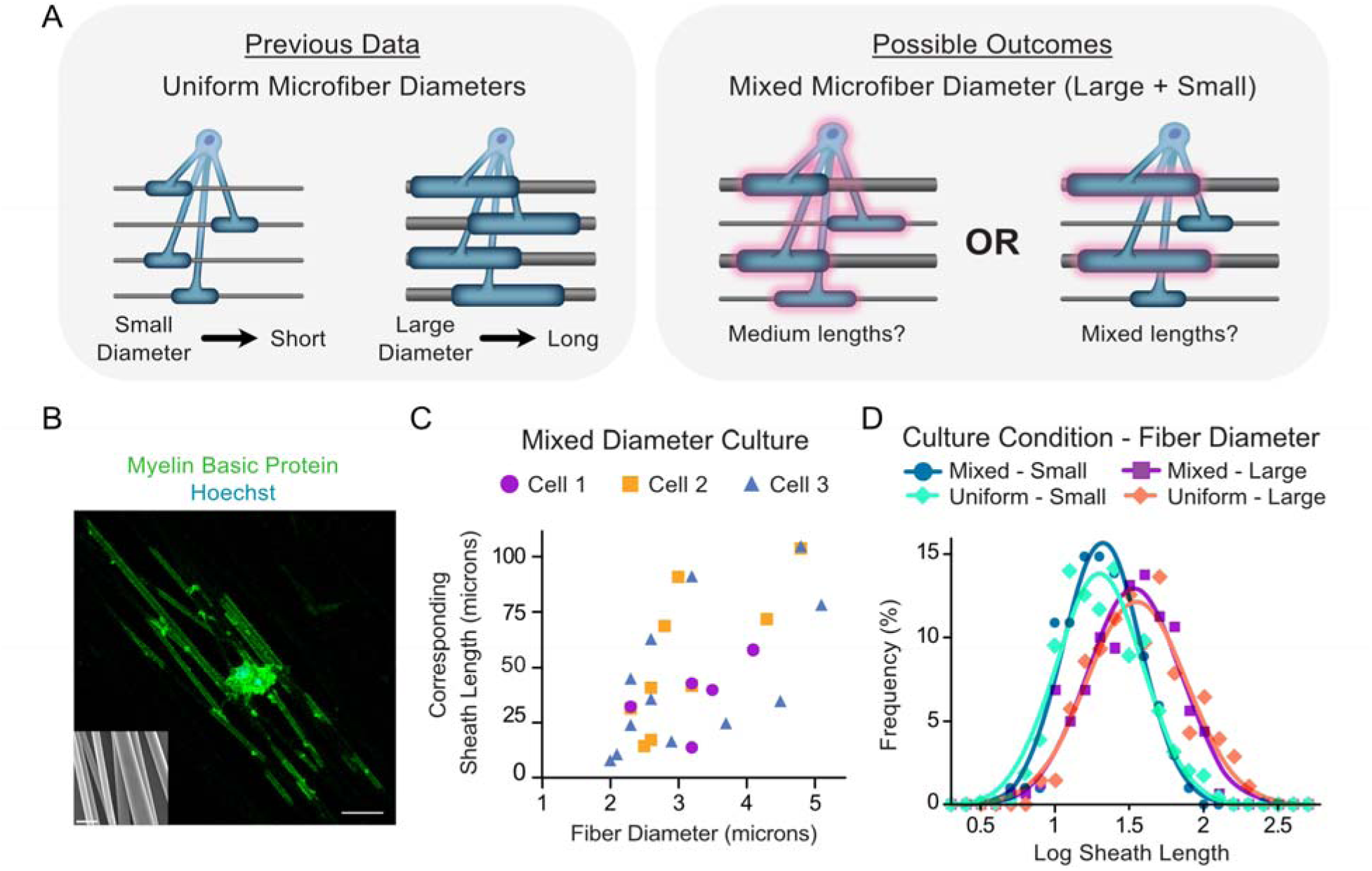
Each nascent myelin sheath locally senses and responds to fiber diameters. **A)** Experiment and predictions. Oligodendrocytes cultured on uniform fiber diameters generate short myelin sheaths on small diameter fibers and long myelin sheaths on large diameter fibers. When oligodendrocytes contact both small and large diameter fibers simultaneously, they will adjust sheath length at the whole cell level (medium lengths) or locally within each nascent myelin sheath (same sheath lengths as they form on uniform fiber diameters). **B)** Confocal maximum projection image of an oligodendrocyte on mixed diameter fibers, forming myelin sheaths on both large and small diameter fibers simultaneously. Scale = 20 µm. SEM inset shows mixed diameter microfibers, scale = 5 µm. **C)** Myelin sheath lengths plotted with the corresponding fiber diameter for three example oligodendrocytes. **D)** Frequency distribution of the log of myelin sheath lengths formed, binned by fiber diameter: small (1 – 2.5 micron) diameter fibers from mixed diameters (solid line, dark blue) or uniform fiber diameter cultures (dashed line, light blue-green), and myelin sheaths formed on large diameter fibers (2.6 – 6 microns) on mixed diameters on (solid line, purple) or uniform (dashed line orange). > 63 individual oligodendrocytes analyzed, from n = 3 independent experiments each with pooled rat litters (>127 sheaths on small diameters, >309 sheaths on large diameters). The distribution of myelin sheath lengths is statistically different between fiber diameters but unchanged between uniform or mixed diameters (t-test, p = 0.42 for small diameter uniform vs mixed, p = 0.21 for large diameter uniform vs mixed, p <0.01 comparing large vs small).

### Piezo1 Enables Local Myelin Sheath Length Scaling

We next set out to address how the diameter is integrated by oligodendrocytes to generate different myelin sheath lengths. When nascent myelin sheaths wrap around axons, myelin membrane curvatures would vary substantially between small diameter (0.5-2.5 microns) and large diameter (2.5-5 microns) axons. Many mechanosensory proteins generate local signaling in response to membrane bending and membrane tension forces; therefore, we cross-referenced mechanosensory proteins in the literature with oligodendrocyte expression profiles^28,38,39^ to generate a list of candidates. We examined the contribution of these candidate proteins to diameter-responsive myelin sheath elongation by utilizing doxycycline-inducible knockdown in primary cortical rat oligodendrocytes cultured on mixed-diameter microfibers. shRNA expression was induced after initial differentiation (based on myelin basic protein (MBP) expression), when initial ensheathment has begun: 4-5 days in culture on microfibers (Figure 2 A). Myelin sheath lengths and the underlying fiber diameters were measured from oligodendrocytes in contact with both large and small diameter microfibers. As expected, control shRNA-expressing oligodendrocytes produced longer myelin sheaths on large diameter microfibers (Figure 2 B,C; Figure S2 A,B) without a substantial impact on the % MBP+ oligodendrocytes (OLs) or OLs ensheathing microfibers (Figure S2 C,D). In contrast, oligodendrocytes expressing shRNA targeting one of our candidates, the mechanosensitive cation channel Piezo1^40^ demonstrated a loss of myelin sheath elongation on the large diameter microfibers (Figure 2 D,E).

**Figure 2.**
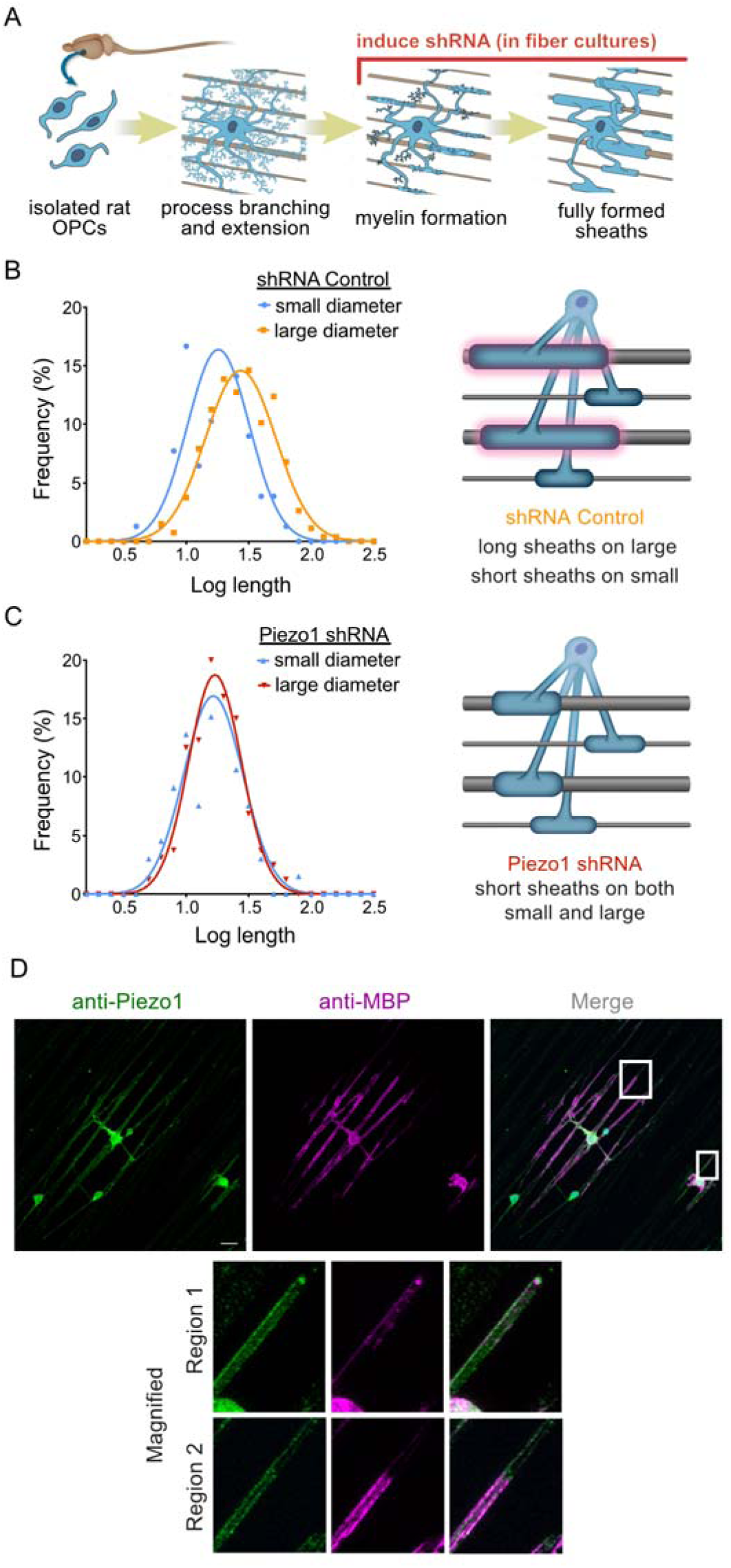
Piezo1 is present in nascent myelin sheaths and is important for myelin sheath elongation on large diameter fibers. **A)** Experimental schematic: shRNA-mediated knockdown of membrane tension-sensing candidate Piezo1 or control shRNA expression was induced after OL differentiation and process extension in mixed diameter fiber cultures. **B)** Log normal curves and schematic representation of myelin sheath length distributions on small or large diameter fibers from mixed diameter cultures with control shRNA. Sheath lengths on large diameter fibers are longer compared to those on small diameter fibers. t-test, p = 0.038. **C)** Log normal curves and schematic of myelin sheath length distributions of sheaths formed on small or large diameter fibers from mixed diameter cultures with Piezo1 shRNA. Sheath length scaling with diameter is lost with Piezo1 shRNA-mediated knockdown. p = 0.64, Welch’s corrected t-test of mean log sheath length on small vs large diameter fibers. n = 4 independent fiber cultures with oligodendrocytes from different pooled rat litters. Error bars = standard deviation. **D)** Maximum projected immunofluorescence image of differentiated oligodendrocytes after 3 days on 2-micron fibers. Magnified regions show localization within and extending beyond forming sheaths. Scale = 20 µm.

In order to locally control sheath length, Piezo1 must be expressed in oligodendrocytes and localize to the myelin sheath. Therefore, we determined the expression and localization of Piezo1. In rats, Piezo1 expression has been shown in oligodendrocyte lineage cells: Piezo1 is present abundantly in oligodendrocyte progenitor cells (Olig2+CC1-) and in mature oligodendrocytes (Olig2+CC1+) at postnatal day 7, during the onset of active developmental myelination^28^. Similarly, we found that Piezo1 mRNA levels were high in acutely isolated platelet derived growth factor receptor alpha (PDGRFα)-positive oligodendrocyte precursors from P6-9 mice. During the latest time points of oligodendrocyte maturation, when myelin oligodendrocyte glycoprotein (MOG) levels and 2′,3′-cyclic nucleotide 3′-phosphodiesterase (CNP) levels rose dramatically, Piezo1 levels began to diminish (Figure S2 E, F). We further assessed whether Piezo1 is localized to nascent myelin sheaths during initiation and elongation by immunostaining rat oligodendrocyte microfiber cultures when the majority of cells expressed MBP and began ensheathment. Consistent with a role in diameter-sensing during initial myelin membrane formation, Piezo1 was found localized in regions of MBP+ membrane ensheathment and at the tips of ensheathment (Figure 2 F and magnified Region 1) as well as in extending processes (Figure 2 F magnified Region 2). Together these data established Piezo1 as a candidate for local control of myelin membrane expansion.

### Piezo1 is Required for Myelin Sheath Length Scaling with Large Diameters In Vivo

To determine whether Piezo1 is critical in vivo for myelin sheath elongation, we knocked out Piezo1 during late differentiation and onset of myelination, using the CNP-Cre line^41^ to drive recombination when crossed with Piezo1 floxed mice^42^. While Piezo1 antibodies work well in rat and human, they have been unsuccessful in mouse unless knocking out both Piezo1 and Piezo2 together^43^. Therefore, we first confirmed loss of Piezo1 expression in isolated mouse oligodendrocytes by mRNA quantification after differentiation. Early differentiation marker CNP mRNA was detected robustly immediately in isolated cells and throughout the culture (Figure S2 F). Reduced Piezo1 levels in conditional knockout mice were confirmed with qPCR after 2 and 4 days of differentiation (Figure 3 A).

**Figure 3.**
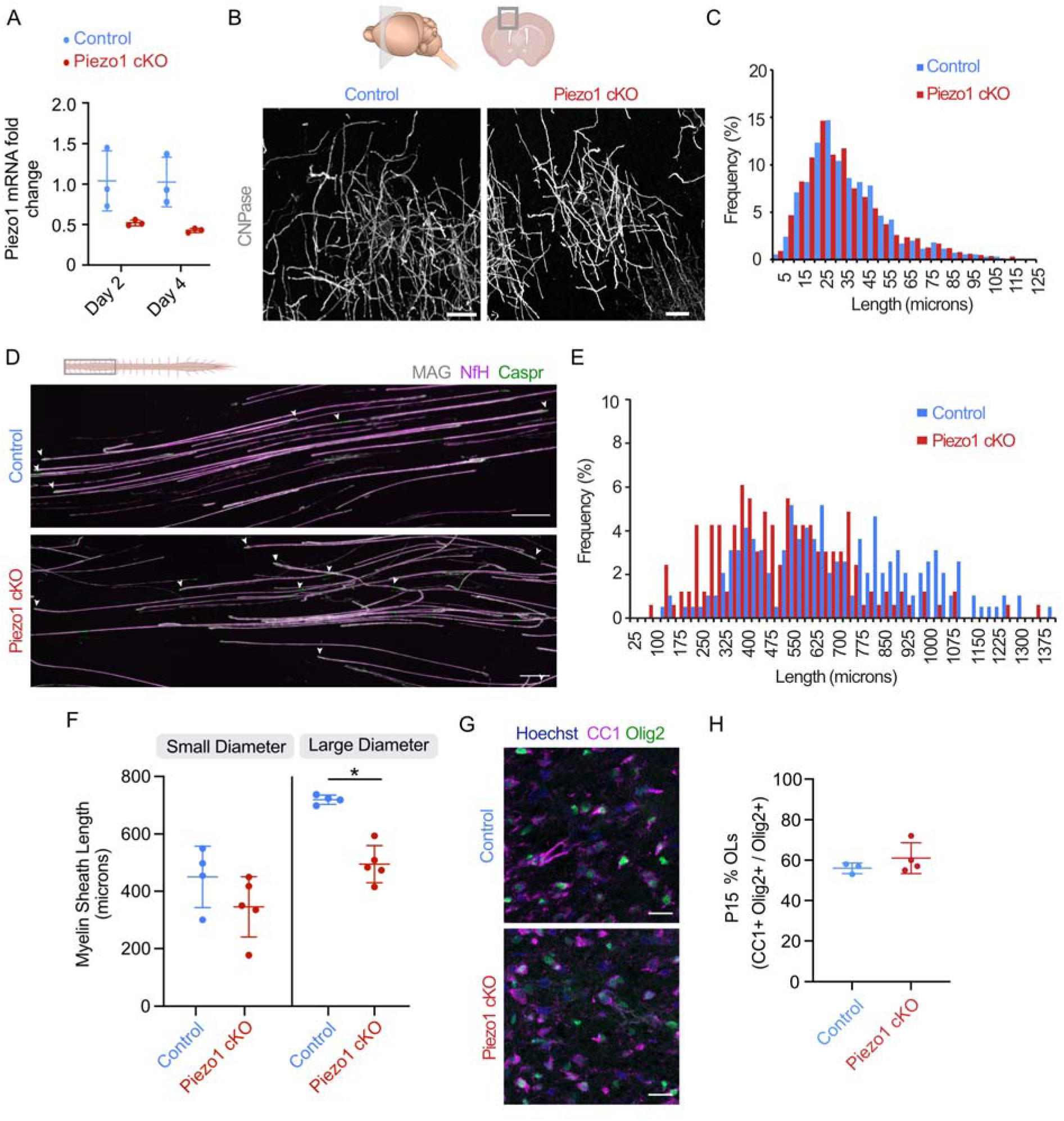
Piezo1 is important for the elongation of myelin sheaths on large diameter axons in vivo. **A)** Relative Piezo1 mRNA levels in isolated mouse OLs during differentiation measured by quantitative PCR. Relative to day 0 of floxed control mice. p = 0.048 between genotypes. **B)** Whole OLs in layers II-III of frontal cortex sections labelled with anti-CNPase. Entire OL cells can be traced to analyze the number and lengths of myelin sheaths formed. **C)** Frequency distribution of myelin sheath lengths formed in the prefrontal cortex binned by 5 microns. **D)** Immunofluorescence of teased spinal cord white matter, labelled for myelin (MAG), axonal (NfH), and paranodal (Caspr) proteins. Ends of myelin sheaths are noted with arrowheads. **E)** Frequency distribution of myelin sheath lengths (25-micron bins) from teased spinal cord axons. 62-142 myelin sheaths measured per mouse from n = 4-5 mice per genotype. **F)** Average myelin sheath lengths on axons smaller than 2.5 microns or greater than 2.5 microns in diameter. p = 0.20 for small diameter, *p = 0.001 for large diameter, Welch’s corrected t-test between genotypes. **G)** Confocal immunofluorescence image of ventral spinal cord white matter at P15, stained for oligodendrocyte lineage marker Olig2 and mature oligodendrocyte marker CC1, alongside Hoechst for nuclei. **H)** Percent of oligodendrocyte lineage cells (Olig2+) that are mature (CC1+Olig2+) at P15. p = 0.39, Welch’s corrected t-test. Error bars = standard deviation.

To establish whether Piezo1 loss impacts the number and lengths of myelin sheaths generated by oligodendrocytes in vivo, we initially assessed oligodendrocytes in the mouse prefrontal cortex at postnatal day 30. Individual oligodendrocytes and all their myelin sheaths can readily be assessed in layers II-III in the mouse cortex at this time point^44^ when peak myelination has occurred, but the area is still undergoing active incorporation of new oligodendrocytes^45,46^. Thick vibratome sections were used to preserve whole oligodendrocytes. Immunostaining for the paranodal marker Caspr alongside CNP and MBP facilitated tracing whole oligodendrocytes and measuring all the connected myelin sheaths (Figures 3 B). Piezo1 cKO mice had both comparable myelin sheath lengths and number of myelin sheaths formed per oligodendrocyte relative to floxed control littermates (Figures 3 C, S3 A-C).

Since the cortex has predominantly small diameter axons (below 2 microns^47^), we turned to the ventral and lateral spinal cord, a region of the mouse CNS with a higher proportion of large diameter axons^48^. To assess myelin sheath lengths in the densely myelinated spinal cord tracts, we used a well-established method of teasing spinal cord white matter in order to distinguish single axons and individual myelin sheaths^49^. Immunostaining for paranodal Caspr, axonal neurofilament H (NfH), and myelin associated glycoprotein (MAG), enabled us to measure individual myelin sheath length and the corresponding axon diameter (Figure 3 D). Piezo1 cKO mice versus the floxed control littermates or CNP-Cre control mice showed substantially reduced myelin sheath lengths (Figures 3 E, S3 D-F). Furthermore, when comparing myelin sheath lengths formed on similar diameter ranges, i.e. small diameters or large diameter axons, Piezo1 cKO reduced the average myelin sheath lengths found on large diameter axons (Figure 3 F). Importantly, since myelin sheath lengths scale with axon diameter in vivo, we verified that the axon diameter distributions were unchanged (Figure S3 G,H). Since CNP is active in differentiating oligodendrocytes, we confirmed that CNP-Cre driven Piezo1 reduction did not impact the differentiation or density of oligodendrocytes by assessing CC1+Olig2+ mature oligodendrocyte at postnatal day 15 in the spinal cord (Figures 3 G,H, S3 I). Altogether, this mirrored our in vitro microfiber results and demonstrated Piezo1 is important in vivo for scaling myelin sheath lengths with large diameter axons.

### Piezo1 is Not Required to Establish Normal Myelin Thickness

Since the number of myelin sheath layers (i.e., thickness) is also known to scale with axon diameter^48^, we set out to determine if Piezo1 is important for this process. To assess the impact on axonal ensheathment and myelin thickness after Piezo1 loss, we performed transmission electron microscopy (TEM) on ultrathin tissue sections from Piezo1 cKO mice at postnatal day 15 and 30. We analyzed the cervical ventral white matter of the spinal cord (Figure 4 A) because of the broad range axon diameters in this region (Figure S4 A,B) and sheath length differences were observed in this region. Consistent with the notion that Piezo1 loss in differentiated oligodendrocytes does not impact the initiation of myelin formation, we found no difference in the percent of myelinated axons at either P15 or P30 (Figure 4 B,C). There was also no difference in the distribution of unmyelinated axon diameters between littermate controls and Piezo1 cKO mice. Unmyelinated axons over 1.5 µm were rare (Figure S4 C-F), demonstrating myelin sheath initiation on large diameter axons is unaffected.

**Figure 4.**
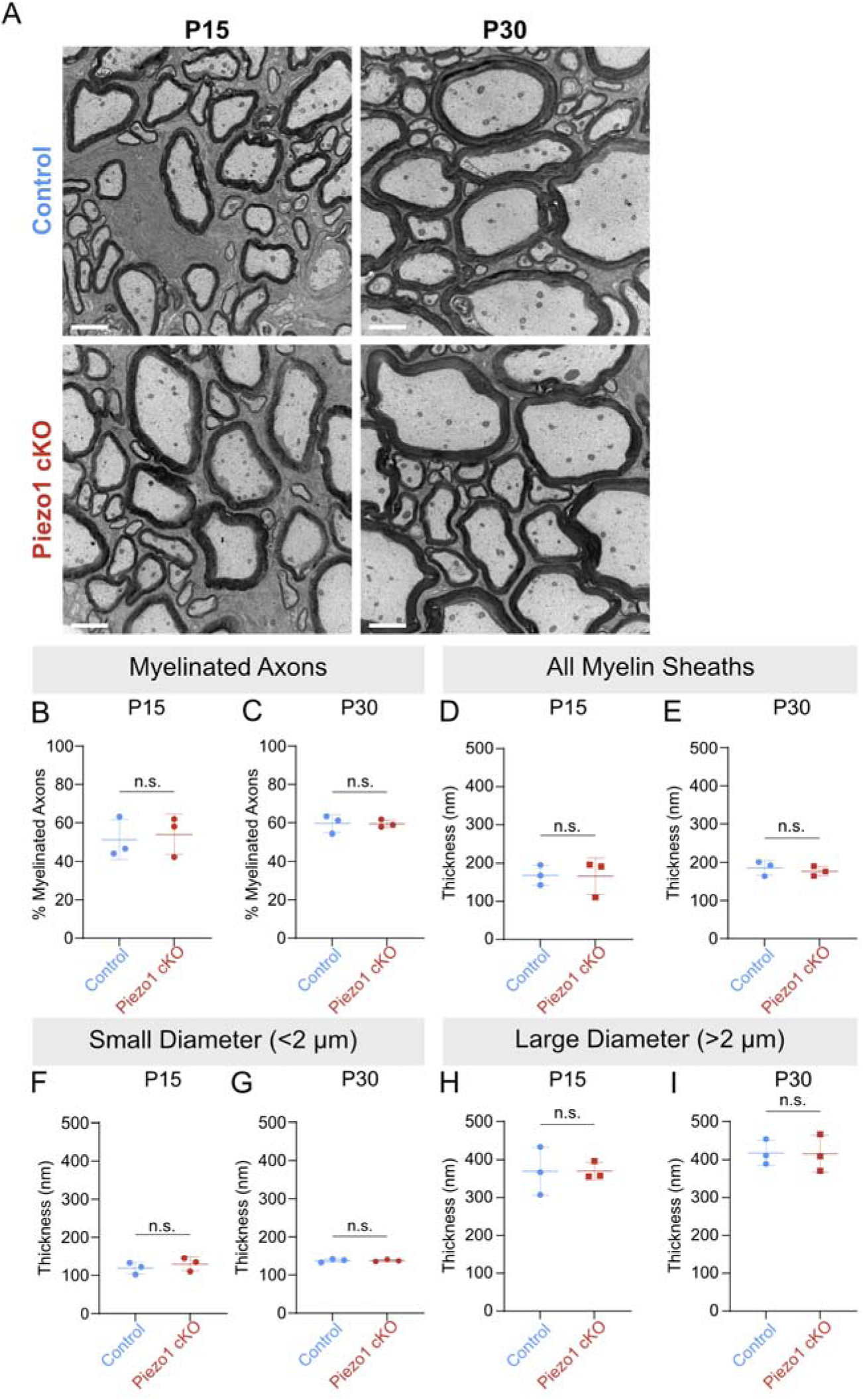
Myelin sheath thickness is not impacted by Piezo1 loss. **A)** Electron microscopy images of ventral cervical spinal cord of control (fl/fl) and Piezo1 cKO mice. Scale = 1 µm. **B, C)** The percent of myelinated axons is unchanged at P15 (p = 0.38) (B) and P30 (p = 0.48) (C). n = 3 mice, >300 axons/mouse. **D-I)** Average myelin sheath thickness at P15 and P30. All p values comparing littermate control (Piezo1 fl/fl) and Piezo1 cKO mice using an unpaired t-test with Welch’s correction. **D)** All myelin sheaths measured at P15, p = 0.47 and **E)** at P30, p = 0.28. n = 3 mice, 189 - 247 sheaths/mouse. **F, G)** Myelin sheath thickness on small diameter axons (< 2 µm) at P15, p = 0.23, and P30, p = 0.49. n = 3 mice, 150 - 212 axons/mouse. **H, I)** Myelin sheaths thickness on the large diameter axons (> 2 µm) at P15, p = 0.49, and P30, p = 0.47. n = 3 mice, 25 - 46 axons/mouse.

Based on the current model of myelin sheath formation suggesting that myelin sheath lengthening is coupled to myelin thickening^50^, we might expect decreased myelin sheath thickness with a myelin sheath length deficit. Interestingly, we found no differences in overall myelin sheath thickness (Figures 4 D,E, S4 G-N). Since Piezo1 impacted sheath lengths in a diameter-dependent manner, we predicted that Piezo1 would impact thickness in a diameter-dependent manner as well. To assess diameter-specific effects, we binned our myelin measurements into groups based on axon diameter. Neither axon diameters less than 2 µm nor greater than 2 µm had differences in myelin thickness (Figure 4 F-I, S4 I-L). Together these results show that Piezo1 does not impact myelin sheath initiation or thickness, regardless of axon diameter.

## Discussion

Myelin sheath size impacts neuronal signal speed^2^. In particular, sheath length is proposed to be instrumental in coordinating signal timing for nervous system function^9,10,11, 51^. If true, understanding how myelin sheath lengths are established and adjust their properties is critical to understanding neural networks and behavior. A combination of adaptive signals (e.g., neuronal activity), an intrinsic program (i.e., oligodendrocyte cell origin), and physical cues (e.g., axon diameter) all contribute. While electrical and biological signaling from other cells certainly can bias axon selection and adjust sheath size during myelin formation^52,53,54,55,56^ studies have shown myelination can occur independently of neuronal activity^18,20,21,22,24^. Importantly, sheath lengths formed independent of neuronal activity are only slightly shorter than those in the presence neuronal activity^52,55^. Furthermore, these active neuronal signals do not explain the striking correlation in the CNS of sheath length and thickness scaling with axon diameter^15,16,48^. Alternatively, we previously proposed that a hard-wired program in oligodendrocytes is used to initiate the myelination process, based on data from neuron-free cultures of oligodendrocytes on inert substrates^25^. Myelin sheath lengths can be established by oligodendrocytes, have intrinsic differences based on oligodendrocyte origin, and adapt sheath lengths in response to physical cues, such as axon diameter^25^. Until now it has remained unclear how ‘hard-wired’ myelination programs respond to axon diameter and if they regulate sheath size globally (cell-wide) or locally, on a sheath-by-sheath basis.

In this work, by using microfibers with varied diameters, we demonstrated that diameter-dependent mechanotransduction controls myelin sheath length locally, within each independent sheath (Figure 1). This is consistent with an earlier electron microscopy observation in mouse tissue where a single oligodendrocyte generated different myelin sheath thicknesses corresponding with axon diameters^36^. Our data provides direct evidence that diameter can control myelin lengths on a sheath-by-sheath level.

Here we identify the Piezo1 mechanosensitive channel as a key protein by which oligodendrocytes detect axon diameter and guide myelin sheath formation locally, in individual sheaths. Knockdown of Piezo1 in oligodendrocytes on mixed-diameter microfibers selectively affected sheath length on large-diameter synthetic axons, whereas sheaths originating from the same oligodendrocytes were unaffected on smaller diameter axons. This differential response suggests that Piezo1 is needed to discern between large- and small-diameter fibers based on physical differences, such as fiber curvature. Piezo1 increases membrane permeability of cations, such as calcium, into cells upon changes in membrane tension and curvature^40,57^. Cryo-EM structural data suggesting that Piezo1 induces local membrane bending in its closed-channel state; in less bent membranes, Piezo1 is flatter and in a more open conformation^58,59^. This is consistent with our data suggesting that less curvature on larger diameter fibers likely shifts the equilibrium of Piezo1 towards the open-channel state.

We propose that the role of Piezo1 in distinguishing large-diameter axons is most important for elongation during early myelin sheath formation. Our results show that Piezo1 is expressed early in differentiated oligodendrocytes then decreases with maturation (Figures 2 F, S2), consistent with reports where Piezo1 expression was observed in OPCs and CC1+ oligodendrocytes in young, but not aged, rats^28^. This also aligns with observations previously reported where calcium transients and waves in myelin sheaths are most prevalent during early myelin formation then decrease over time^60,61^. Thus, we hypothesize Piezo1 contributes to these early calcium dynamics to establish diameter-dependent sheath lengths. This agrees with Auer et al., who demonstrated that in vivo myelin sheath length differences are established within a 3-day window from initiation. During this early time window, myelin sheaths undergo their most rapid elongation^62^, paralleling the observed expression profile of Piezo1.

Strikingly, our in vivo data demonstrates differential regulation of myelin sheath length and thickness (number of layers around an axon). Sheath lengths are smaller in Piezo1 cKO teased spinal cord preparations compared to controls (Figures 3, S3) without any significant changes to sheath thickness (Figures 4, S4). This data challenges the idea of inherent coupling of sheath length and thickness. Scaling of both myelin sheath length and thickness with axon diameter has been described since the late 1940s^15,16,17,48^. Over the years, many models of myelination have been proposed but have kept these two parameters intertwined. For example, the model proposed by Snaidero et al. in 2014 suggests that spiral wrapping of myelin around the axon drives lateral elongation of myelin sheaths^50^. Our data strongly supports that the mechanisms regulating myelin longitudinal growth and membrane wrapping (thickness) are independent pathways. Other studies also point to independent regulation of myelin sheath length and thickness. For example, monocular deprivation has been shown to decrease sheath length, without impacting overall sheath thickness^52^. Small reductions in MBP mRNA have been shown to decrease sheath thickness but not sheath length^63^. Further, genetic mouse models that increase myelin outfoldings and swellings, such as oligodendrocyte-specific dampening of calcium signaling or loss of ubiquitin ligase Fbxw7, show consequences on overall myelin sheath length but not thickness^64,65^. In particular, the finding that dampening calcium in oligodendrocytes affects myelin sheath length with little evidence of an effect on thickness^64^ is of special interest due to the tentative connection of Piezo1 to oligodendrocyte calcium signaling, as a cation channel. Together, this supports a connection between calcium signaling and myelin sheath lengthening but not sheath thickness during developmental myelination.

It has been proposed that even small changes in myelin morphology affect axonal conduction, which can impact the precise temporal control of neuronal firing^8,52,66^. Indeed, some studies have proposed that the property of myelin sheath length could be the primary means to coordinate timing in particular neural circtuits^7,8^. Direct experimental evidence demonstrating the impact on neuronal signaling when only CNS myelin sheath lengths change (e.g., without concomitant changes to thickness) are lacking. Piezo1 cKO mice display an average of 225 microns shorter myelin sheaths on large-diameter axons (Figure 3) without thickness differences. Thus, this work highlights a potential future avenue to experimentally uncouple the impact of length and thickness on neuronal signaling.

In this work, we address a longstanding question of how oligodendrocytes sense and scale myelin sheath lengths with axon diameter. This correlation has implications for neuronal signaling and has been observed in vivo for over 75 years without a mechanism. Our data points to a model in which oligodendrocytes locally sense axon diameter in the earliest stages of myelination through Piezo1, during a period of rapid myelin sheath elongation. Our data supports this model by demonstrating myelin sheath length is regulated at the individual myelin sheath level and by showing Piezo1 plays a critical role in myelin sheath elongation on large diameter axons. We also show that myelin sheath length and thickness can be uncoupled, as reduction in sheath elongation did not impact myelin sheath layers. This challenges existing models of myelination by demonstrating independent regulation of length and thickness and serves to further our understanding of mechanisms underlying myelination during central nervous system development.

## Materials and Methods

All animal procedures were conducted in accordance with procedures approved by the SUNY Upstate Medical University’s institutional animal care and use committee.

### Primary Rat Oligodendrocytes

Oligodendrocyte precursor cells (OPCs) were isolated from P0-P2 Sprague-Dawley rats (Charles River). Cerebral cortices were collected and pooled from one or more litters. Meninges were removed from isolated cortices, then cortices were minced and enzymatically dissociated for 1 h at 37°C with 1.2 U/mL papain (Worthington), 0.1mg/mL L-cysteine (MilliporeSigma) and 0.40 mg/mL DNase I (MilliporeSigma). Mixed glial cells were plated at a density of 1.5 cortices per poly-D-lysine (PDL) coated T75 flask and cultured at 37°C in 7.5% CO_2_ in high-glucose DMEM (Gibco), 10% fetal bovine serum (FBS, Gibco and MilliporeSigma) and 1% penicillin/streptomycin (pen/strep, Gibco). After 10 days, OPCs were enriched by mechanical shaking^67^. Shake-off cells were plated onto petri dishes for 30 m to further remove unwanted glia and non-adherent OPCs were collected for culturing.

### Mice

Piezo1 flox/flox mice, with LoxP sites flanking exons 20-23, were originally generated by the Patapoutian lab^42^ and were obtained from Jackson Laboratories (stock# 029213). To generate mice with conditional Piezo1 knockout in differentiating cells (cKO), homozygous Piezo1^fl/fl^ animals were crossed with mice with heterozygous expression Cre recombinase under the control of the 2′ 3′ cyclic nucleotide phosphodiesterase (CNP) gene^41^ to generate flox control (CNP^WT/WT^; Piezo1^fl/fl^) and cKO littermates (CNP^WT/Cre^; Piezo1^fl/fl^). CNP-Cre control mice (CNP^WT/Cre^) were sourced from breeding CNP^WT/WT^ with CNP^WT/Cre^ parents. Mice were either genotyped by Transnetyx or using the following primers. CNP-Cre primers:

CATAGCCTGAAGAACGAGA, GATGGGGCTTACTCTTGC, GCCTTCAAACTGTCCATCTC; Piezo1 flox primers: GCC TAG ATT CAC CTG GCT and GCT CTT AAC CAT TGA GCC ATC T.

### Primary Mouse Oligodendrocytes

Immunopanning of mouse OPCs from P6-P9 pup cerebral cortices was done by a modified method based upon^67^. After euthanasia and collecting cerebral cortices, a tail clip was taken for subsequent genotyping. Each pair of cerebral cortices was enzymatically and mechanically dissociated using a gentleMACS dissociator and the Neural Tissue Dissociation Kit P (Miltenyi). Cells were resuspended to single cell suspension in 0.2% BSA and 5 µg/mL insulin in Dulbelco’s PBS (MilliporeSigma), then negatively selected with Griffonia Simplicifolia Lectin (Vector Labs)-coated petri dishes twice for 15 m. Cell suspensions were then transferred to anti-PDGFRα (CD140a, BioLegend Cat# 135902, RRID:AB_1953328) coated petri dishes for 45 m for positive selection. Attached cells were collected. Cells were pelleted and resuspended prior to plating on a PDL-coated cell culture dish in culture medium. Medium was composed of DMEM:Neurobasal Media, B-27 alternative made in-house (as below), 5 μg/mL N-acetyl cysteine, and 10 ng/mL D-biotin, ITS supplement (MilliporeSigma), and modified Sato (100 µg/mL BSA fraction V, 60 ng/ml Progesterone, 16 µg/ml Putrescine, 40 ng/mL Tri-iodothyroxine; reagents from MilliporeSigma) and penicillin–streptomycin. The house-made B-27 (reagents from MilliporeSigma) was based upon Chen et al. 2008^68^, with modifications as follows: bovine serum albumin (Cat# A1470), synthetic L-carnitine hydrochloride (Cat# C0283), and synthetic (±)-α-Tocopherol (Cat# T3251). Media was supplemented with growth factors 10 ng/ml PDGF-AA, 5 ng/ml neurotrophin 3 (NT3) (Millipore Sigma) and 10 ng/ml ciliary neurotrophic factor (CNTF) (Peprotech) and 2.05 µg/mL forskolin (Cayman Chemical Cat# 11018). Cells were cultured at 37°C and 7.5% CO_2_ with a half media change every other day with new growth factors.

### qPCR

Piezo1 expression analysis and confirmation of knockout in oligodendrocytes was performed by qPCR. Oligodendrocytes were isolated from CNP+/Cre, CNP +/Cre; Piezo1 fl/fl, and Piezo1 fl/fl mice. Isolated mouse OPCs were divided evenly between Trizol lysis (for Day 0) and PDL-coated cell culture plates for collection on days 2, 4, 6, 8, and/or 10. RNA was extracted from cells with Trizol followed by chloroform extraction using a Monarch Total RNA Miniprep Kit. DNase IXT (New England Biolabs) was used for in-column DNase treatment. cDNA generated with the LunaScript RT SuperMix Kit (New England Biolabs). IDT PrimeTime Standard qPCR assay and qPCR probe and primer sets were used for qPCR. Primer and probe sets were as follows:

CNP: AGAGAGCAGAGATGGACAGT, AATTCTGTGACTACGGGAAGG,
/5Cy5/AGCAGGAGG/TAO/TGGTGAAGAGATCGTA/3IAbRQSp/
Piezo1: CATGCGTTGCCACTCCT, GGCTGTACCTACCTGACTTCT,
/5SUN/CCTTATCAG/ZEN/TGACTTCCTCCTGCTGC/3IABkFQ/
GAPDH: GTGGAGTCATACTGGAACATGTAG, AATGGTGAAGGTCGGTGTG, /56-FAM/TGCAAATGG/ZEN/CAGCCCTGGTG/3IABkFQ/
MOG: AGTCCGATGGAGATTCTCTACT, CACTTGTGCCTACGATCCTC,
/5Cy5/CACGAAGTT/TAO/TTCCTCTCAGTCTGTGCT/3IAbRQSp/

Multiplexed qPCR was performed on a BioRad CFX Opus. All Ct values were normalized against GAPDH values, to obtain ΔCt. For expression across time, ΔΔCt was normalized against expression at time of cell isolation. Fold change expression between Piezo1 cKO and floxed control mouse cells were calculated with 2 ^-(ΔΔCt)^ relative to floxed controls at Day 0.

### Electrospun Fiber Cultures

Electrospun microfibers composed of poly-L-lactic acid were synthesized as custom, parallel-aligned fibers and suspended by fitting into 12-well plate inserts by The Electrospinning Company. Specific fiber diameter ranges of 1-2 µm, and 2-4 µm were used as reported previously^24^. Mixed fiber diameter microfiber scaffolds spanning fiber diameters between 0.7 - 6 microns distributed across each scaffold were generated and verified with scanning electron microscopy by The Electrospinning Company. Microfiber scaffolds were soaked in 70% ethanol for 10 m, washed, then coated with PDL. Rat OPCs were added at 30,000-35,000 per scaffold in myelin medium^35^. Medium was changed every 2-3 days. Myelin medium was as follows: 50:50 high-glucose DMEM:Neurobasal Media, B27 (Gibco) or a B-27 alternative (as above), 5 μg/mL N-acetyl cysteine, and 10 ng/mL D-biotin, ITS supplement (MilliporeSigma), and modified Sato (as above but with 400 ng/mL Tri-iodothyroxine, 400 ng/mL L-Thyroxine; reagents from MilliporeSigma), and penicillin–streptomycin.

### Inducible Knockdown of Gene Expression

Rat OPCs were transduced with SMARTvector inducible lentiviruses (TU at least 10^7^/mL) for shRNA-mediated knockdown generated by Horizon Discovery. shRNA sequences used are as follows: Piezo1 shRNA: 1-AAGTACGACCTGGTGCAAC, 2 – TCACGGGCATCTACGTCAA, 3 – TGCTGTGCCTCACGTGTT. These Tet-On®3G inducible viruses were generated to express shRNA targets using mouse CMV promoter, including turboGFP as a reporter of expression. Non-targeting control shRNA (VSC6584, HorizonDiscovery) was used to account for non-specific effects of lentiviral shRNA and turboGFP. Pooled lentiviruses generated against three target shRNA sequences were used. Prior to lentiviral transduction, 200,000 rat OPCs were plated into each well of a PDL-coated 6-well plate with proliferation medium. After 24 h cells were given fresh proliferation medium and incubated overnight with lentivirus at 5 MOI, as optimized based upon turboEGFP expression and cell viability after 72 h. Proliferation medium was as follows: high-glucose DMEM (Gibco), ITS (Sigma), and modified Sato (as above), pen–strep, and 0.5% FBS with fresh 10 ng/ml PDGFa and 10 ng/ml bFGF (Peprotech). After overnight incubation with viruses, cells were washed with D-PBS (Sigma), dissociated from the plates with TrypLE (Gibco), spun down and resuspended in myelin medium, then plated at 35,000 OPCs per 12-well microfiber insert.

### Microfiber Culture Immunofluorescence and Analysis

After 14 days in culture, cells were fixed in 4% formaldehyde/PBS, followed by PBS washes, and permeabilization in 0.1% TritonX-100 in PBS. Primary antibodies were diluted in PBS and incubated overnight at 4°C. Primary antibodies were: anti-rat myelin basic protein 1:250 (Biorad MCA409S, RRID: AB_325004), chicken anti-GFP 1:500 (Abcam ab13970, RRID: AB_300798). Cells were washed in PBS and incubated 1 h with Alexafluor 488, 568, or 647 conjugated secondary antibodies (Invitrogen), used at 1:1000. After three PBS washes, cells were stained with 5 µg/mL Hoechst (Sigma-Aldrich), washed again and mounted with Fluoromount G (Southern Biotech) with cover glass onto glass slides.

Images were obtained on a Leica SP8 confocal scanning microscope, with a 40x oil/NA1.3 objective. Confocal stacks of 0.35 µm z-steps were taken at 1024 x 1024. At least 10 random areas containing MBP positive cells were imaged across each coverslip. All settings were kept the same within each replicate experiment and images were blinded.

Fiji ImageJ was used to analyze myelination. For comparisons of myelin sheaths on mixed diameter fibers, it was not possible to blind the experimenter from the fiber diameter. Sheaths were defined as > 4.5 micron-long continuous MBP positive membranes fully surrounding a microfiber as assessed using the 0.35 µm z-series. Tubes were traced to measure the length and a perpendicular line was drawn to measure the corresponding fiber diameter. The frequency of sheath lengths was calculated for 5 µm bins. Frequencies from at least three independent experiments were generated and plotted as a frequency distribution. The sheath lengths were determined to be log gaussian distributions, therefore mean log lengths were used for statistics. Binned log lengths (0.1 bin size) were also used to generate frequency distribution plots. For each oligodendrocyte analyzed, myelin sheaths formed on different caliber fibers were measured from the same cell. At least three independent experiments were cells pooled from different rat litters on different days. A minimum number of sheaths were measured for microfiber experiments, as indicated in figure legends.

### CNS Tissue Acquisition for Teased Spinal Cords, Vibratome and Cryosections

Mice were euthanized by cardiac perfusion fixation under anesthesia by flushing with PBS followed by 4% formaldehyde in PBS. Brain and spinal cord tissue was isolated immediately after formaldehyde perfusion fixation of mice. Tissue was post-fixed in 4% formaldehyde in PBS either for 30 m or overnight at 4°C (see below).

### Immunofluorescence of Tissue Cryosections and Differentiation Analysis

Overnight post-fixed and PBS washed spinal cord tissue was cryoprotected with 30% sucrose, then flash frozen in dry ice-cooled 2-methylbutane and stored at −80°C. Cervical spinal cords were cut coronally with a Leica cryostat to generate 10-15 μm sections mounted onto Superfrost Plus slides (Fisherbrand). Cryosections were blocked in 10% goat serum and 0.1% TritonX-100 in PBS. Primary antibodies were diluted in blocking solution and incubated overnight at 4°C. Primary antibodies were: anti-CC1 1:500 (Abcam Ab16794, RRID: AB_443473), anti-Olig2 1:500 (RnD AF2418, RRID: AB_2157554) and anti-MBP 1:500 (BioRad MCA409S RRID: AB_32500). Sections were washed with PBS, followed by species-specific Alexafluor 488, 555, or 633 antibodies 1:1000 in blocking solution for 1 h. Cryosections were washed with PBS, stained with Hoechst, and mounted with Fluoromount G.

Images were obtained on a Leica SP8 confocal scanning microscope with a 40x water/NA1.1 objective. Stacks of 1 µm z-steps were taken at 1024 x 1024. At least two sections with a minimum of 60 µm spacing were imaged. For each tissue section, four images were acquired (two lateral, two ventral white matter). The same settings were universally applied within each replicate experiment and images were blinded. Cells that were positive for Hoechst, Olig2, and CC1 were counted from max projected images in Fiji ImageJ, using synchronize windows and the multi-point tool. Hoechst, CC1 and Olig2 positive cells from the total number Hoechst and Olig2 positive cells were calculated to assess % differentiated oligodendrocytes.

### Immunofluorescence and Analysis of Cortical Brain Sections

Cortical brain tissue preparation and analysis was done with minor modifications as in^44,63^. Overnight post-fixed and PBS washed tissue was embedded in 2% low melting point agarose in PBS. The medial prefrontal cortex was sectioned coronally in 150 μm slices on a Leica VT1000 vibratome. The prefrontal cortex slices used were between bregma 1.3 mm and 1.9 mm in the infralimbic and prelimbic areas. Free-floating sections were processed for antigen retrieval by 20 m incubation at 95°C in 0.05% Tween20, 10 mM tri-sodium citrate (pH 6.0). Sections were blocked for 3 h in 10% goat serum and 0.25% TritonX-100 in PBS. Primary antibodies, diluted in blocking solution, were incubated rocking at 4°C for 24 h. Primary antibodies were: anti-CNPase 1:2000 (Sigma C5922, RRID: AB_476854), anti-MBP 1:250 (BioRad MCA409S RRID: AB_32500), anti-Caspr 1:250 (Abcam ab34151, RRID: :AB_869934). Sections were washed in PBS followed by 4 h incubation with 1:1000 diluted Alexafluor 488, 568, or 647 conjugated antibodies in blocking solution. Nuclei were stained with Hoechst followed by mounting sections onto slides with Fluoromount G.

Images were obtained on a Leica SP8 confocal scanning microscope, with 40x oil/NA1.3, and 40x oil/NA1.25 objectives. Stacks of 1 µm z-steps were taken at 1024 x 1024 resolution. Eight random fields of view (290 µm x 290 µm) of the medial prefrontal cortex layer II/III were imaged per mouse. Settings were kept the same across all brain slice images, and images were blinded.

Three-dimensional analysis of myelin sheath lengths was performed using the simple neurite tracer plugin^69^. Myelin sheaths were analyzed for CNP and MBP positive oligodendrocytes. Only myelin sheaths and cells where all sheaths were contained within the stack were quantified. This was assessed by following each CNP positive process from the cell body through the entire z-stack. The lengths of CNP-positive segments flanked by paranodal Caspr bands were measured. The sheath lengths were log normal, therefore both the frequency of binned raw sheath lengths as well as binned log sheath lengths were used. For statistics, a two-tailed t-test with Welch’s correction was used to compare the average log sheath length, with individual mice as the biological replicates. Sample sizes were based on prior publications and power calculations, with power 0.8, alpha 0.05, and a 10-micron effect size^44,63^.

### Teased Spinal Cord Preparation and Immunofluorescence

Cervical and thoracic spinal cord regions from tissue post-fixed 30 m, followed by short-term storage up to 48 h in PBS at 4°C was used. Strips of ventral white matter were teased onto SuperFrost Plus slides with acupuncture needles and immunostained following established procedures^49^. Briefly, after 1 h of permeabilization with 3% normal donkey serum, 2% bovine serum albumin, 0.1% Triton X-100 in PBS, primary antibodies were diluted in this buffer overnight. Primary antibodies were: neurofilament (NfH) 1:1000 (BioLegend 822601, RRID:AB_2564859), Caspr 1:500, MAG 1:100 (Santa Cruz Biotechnology sc-9544, RRID:AB_670102 and R and D Systems AF538, RRID:AB_355423). After washes, secondary AlexaFluor antibodies were incubated in the same buffer for 1h, followed by additional washes and mounting with FluoromountG and cover glass.

Images were obtained on a Nikon Eclipse Ti2 with a Plan Apo 20x/NA0.8 objective. Random areas were imaged as tiled 0.5 µm z-step stacks at 2048 x 2048. Tiled images were stitched with the Image J pairwise or grid collection stitching plugin^70^ to assure inclusion of sheaths greater than 1000 µm in length. Myelin sheath lengths were measured along NfH and MAG positive myelinated axons from paranode to paranode (Caspr band to Caspr band). The axon diameters were assessed by perpendicular measurements acquired at approximately 10 µm from the paranodes. Myelin sheath lengths were both averaged per mouse as well as plotted as frequency of 25 µm binned lengths. The average and distribution of lengths were also binned into sheaths formed on axons above or below 2.5 µm in diameter. Statistics were conducted on average lengths, with individual mice considered the biological replicate. Sample sizes were based on prior publications and power calculations, with power 0.8, alpha 0.05, and a 120 µm effect size^49,63^.

### Transmission Electron Microscopy

Mice were anesthetized and cardiac perfusion fixed at postnatal days 15 and 30. Mice were perfused with 0.1 M phosphate buffer for 2-3 m then a 4% PFA/2.5% Glutaraldehyde mix in 0.1 M phosphate buffer for 10 m at 1 mL/min. The spinal cord was dissected out, meninges removed, sub-dissected into 1 mm pieces, then post-fixed overnight at 4°C. Tissue was washed in phosphate buffer, further fixed in 1% OsO_4_ with 1.5% potassium ferrocyanide, then dehydrated with 10-minute ethanol wash series (25% x1, 50% x2, 70% 2x, 90% 2x, 100% x4). Samples were rinsed with acetone, followed by infiltration with a series of transitional solvent/Epon 812 mixtures on a rotator, as follows: 2 h in a 2:1 acetone:Epon 812 mix, overnight in 1:1, 2 h in 1:3 acetone:Epon 812, then pure Epon 812 four times for a minimum of 1 h each. Sample were cured at 60°C. Ultrathin sectioning and TEM were performed at the SUNY Upstate Medical University TEM Core. Cervical spinal cord samples were cut into 70-90 nm sections using a Leica microsystems EM UC7 ultramicrotome and Diatome diamond knife. Sections were post-stained in Uranyless (Electron Microscopy Sciences), washed in distilled water, Lead citrate stained (Electron Microscopy Sciences) and washed again in distilled water. Ventral white matter images adjacent to the median fissure were imaged with a Jeol JEM-1400 series 120 kV TEM operated at 80 kV and a Gatan Orius SC-1000 CCD camera.

### Transmission Electron Microscopy Analysis

Quantification was performed blinded using FIJI/ImageJ. Axon diameter was determined by measuring the perimeter of the axon, then dividing by pi. Myelin sheath thickness was determined by averaging 5-10 measurements of the electron dense compact myelin layers, excluding any non-compacted areas (e.g., the inner tongue or splitting in the layers). Axons in the ventral white matter of the cervical spinal cord were measured from 4-5 non-adjacent fields of view, yielding ∼200 myelinated axons/mouse. For percentage of myelinated axons, axons from the 4-5 fields of view were categorized as either unmyelinated or myelinated, which includes ensheathed axons without electron dense compacted myelin (>300 axons/mouse). Replicates were individual mice, with sample size based on power calculations power > 0.8, alpha 0.05, for and effect size of 50 – 70 nm for all axons.

### Statistics

Error bars presented are the standard deviation to show the variation. Statistical analysis was done in GraphPad Prism. Data was tested for normality using a Shapiro Wilks test. For two groups, unpaired two-tailed t-Tests with Welch’s correction were used. For comparisons of 3 groups, one-way Anova was used. Biological replicates and sample size was determined prior to experimentation using G*power with anticipated effect sizes from data generated in prior publications, as described in each section above.

## Acknowledgements

We would like to thank Dr. Matt Swire and Dr. Andrew Jarjour for advice and initial assistance with cortical myelination and spinal cord teased fiber preparations, respectively. We thank Benjamin Zink in the SUNY Upstate Medical University Electron Microscopy core for preparation of ultrathin sections and imaging assistance. Additionally, we would like to thank the Centre of Regenerative Medicine at the University of Edinburgh where work was initiated, as well as past members of the ffrench-Constant lab, Williams lab, and Veronique Miron for helpful discussions. Support for this research was initially provided by the Wellcome Trust (Investigator Award to CffC) and then provided by the Esther A & Joseph Klingenstein Fund, the Simons Foundation, and National Institutes of Health National Institute of Neurological Disorders and Stroke R01NS135206 to MEB.

## Author Contributions

**ARY-** Conceptualization, Formal analysis, Experimentation, Visualization, Supervision, Writing – original draft, Writing – review and editing; **RWL-** Conceptualization, Experimentation, Formal analysis, Visualization, Writing – review and editing**; MC –** Experimentation, Writing – editing**; BP-** Analysis**; MZ-** Analysis**; JR –** Validation, Experimentation, Analysis**; AD –** Experimentation and Analysis**; MEB –** Conceptualization, Methodology, Experimentation, Formal analysis, Writing – original draft, Writing – review and editing, Visualization, Supervision, Project administration, Funding acquisition

## References

1. Simons, M., and Nave, K.-A. (2016). Oligodendrocytes: Myelination and axonal support. Cold Spring Harb Perspect Biol 8, a020479. 10.1101/cshperspect.a020479.

2. Brill, M.H., Waxman, S.G., Moore, J.W., and Joyner, R.W. (1977). Conduction velocity and spike configuration in myelinated fibres: computed dependence on internode distance. J Neurol Neurosurg Psychiatry 40, 769–774. 10.1136/jnnp.40.8.769.

3. Waxman, S.G. (1980). Determinants of conduction velocity in myelinated nerve fibers. Muscle Nerve 3, 141–150. 10.1002/mus.880030207.

4. Butt, A.M., Ibrahim, M., and Berry, M. (1998). Axon-myelin sheath relations of oligodendrocyte unit phenotypes in the adult rat anterior medullary velum. J Neurocytol 27, 259–269.

5. Murtie, J.C., Macklin, W.B., and Corfas, G. (2007). Morphometric analysis of oligodendrocytes in the adult mouse frontal cortex. J Neurosci Res 85, 2080–2086. 10.1002/jnr.21339.

6. Chong, S.Y.C., Rosenberg, S.S., Fancy, S.P.J., Zhao, C., Shen, Y.A.A., Hahn, A.T., McGee, A.W., Xu, X., Zheng, B., Zhang, L.I., et al. (2012). Neurite outgrowth inhibitor Nogo-A establishes spatial segregation and extent of oligodendrocyte myelination. Proc Natl Acad Sci U S A 109, 1299–1304. 10.1073/pnas.1113540109.

7. Seidl, A.H., Rubel, E.W., and Harris, D.M. (2010). Mechanisms for adjusting interaural time differences to achieve binaural coincidence detection. J Neurosci 30, 70–80. 10.1523/JNEUROSCI.3464-09.2010.

8. Ford, M.C., Alexandrova, O., Cossell, L., Stange-Marten, A., Sinclair, J., Kopp-Scheinpflug, C., Pecka, M., Attwell, D., and Grothe, B. (2015). Tuning of Ranvier node and internode properties in myelinated axons to adjust action potential timing. Nat Commun 6, 8073. 10.1038/ncomms9073.

9. Stanford, L.R. (1987). Conduction velocity variations minimize conduction time differences among retinal ganglion cell axons. Science (1979) 238, 358–360. 10.1126/science.3659918.

10. Waxman, S.G. (1997). Axon-glia interactions: Building a smart nerve fiber. Curr Biol 7, R406–R410. 10.1016/S0960-9822(06)00203-X.

11. Fields, R.D. (2005). Myelination: An overlooked mechanism of synaptic plasticity? The Neuroscientist 11, 528–531. 10.1177/1073858405282304.

12. Vasistha, N.A., Johnstone, M., Barton, S.K., Mayerl, S.E., Thangaraj Selvaraj, B., Thomson, P.A., Dando, O., Grünewald, E., Alloza, C., Bastin, M.E., et al. (2019). Familial t(1;11) translocation is associated with disruption of white matter structural integrity and oligodendrocyte–myelin dysfunction. Mol Psychiatry 24, 1641–1654. 10.1038/s41380-019-0505-2.

13. Khelfaoui, H., Ibaceta-Gonzalez, C., and Angulo, M.C. (2024). Functional myelin in cognition and neurodevelopmental disorders. Cell Mol Life Sci 81, 181. 10.1007/s00018-024-05222-2.

14. Prineas, J.W., and Connell, F. (1979). Remyelination in multiple sclerosis. Ann Neurol 5, 22–31. 10.1002/ana.410050105.

15. Hess, A., and Young, J.Z. (1949). Correlation of internodal length and fibre diameter in the central nervous system. Nature 164, 490–491. 10.1038/164490a0.

16. Murray, J.A., and Blakemore, W.F. (1980). The relationship between internodal length and fibre diameter in the spinal cord of the cat. J Neurol Sci 45, 29–41. 10.1016/S0022-510X(80)80004-9.

17. Ibrahim, M., Butt, A.M., and Berry, M. (1995). Relationship between myelin sheath diameter and internodal length in axons of the anterior medullary velum of the adult rat. J Neurol Sci 133, 119–127. 10.1016/0022-510X(95)00174-Z.

18. Lee, S., Leach, M.K., Redmond, S.A., Chong, S.Y.C., Mellon, S.H., Tuck, S.J., Feng, Z.Q., Corey, J.M., and Chan, J.R. (2012). A culture system to study oligodendrocyte myelination processes using engineered nanofibers. Nat Methods 9, 917–922. 10.1038/nmeth.2105.

19. Goebbels, S., Wieser, G.L., Pieper, A., Spitzer, S., Weege, B., Yan, K., Edgar, J.M., Yagensky, O., Wichert, S.P., Agarwal, A., et al. (2017). A neuronal PI(3,4,5)P3-dependent program of oligodendrocyte precursor recruitment and myelination. Nat Neurosci 20, 10–15. 10.1038/nn.4425.

20. Rosenberg, S.S., Kelland, E.E., Tokar, E., De La Torre, A.R., and Chan, J.R. (2008). The geometric and spatial constraints of the microenvironment induce oligodendrocyte differentiation. Proc Natl Acad Sci 105, 14662–14667. 10.1073/pnas.0805640105.

21. Lundgaard, I., Luzhynskaya, A., Stockley, J.H., Wang, Z., Evans, K.A., Swire, M., Volbracht, K., Gautier, H.O.B., Franklin, R.J.M., ffrench-Constant, C., et al. (2013). Neuregulin and BDNF induce a switch to NMDA receptor-dependent myelination by oligodendrocytes. PLoS Biol 11, e1001743. 10.1371/journal.pbio.1001743.

22. Koudelka, S., Voas, M.G.G., Almeida, R.G.G., Baraban, M., Soetaert, J., Meyer, M.P.P., Talbot, W.S.S., and Lyons, D.A.A. (2016). Individual neuronal subtypes exhibit diversity in CNS myelination mediated by synaptic vesicle release. Curr Biol 26, 1447–1455. 10.1016/j.cub.2016.03.070.

23. Mayoral, S.R., Etxeberria, A., Shen, Y.-A.A., and Chan, J.R. (2018). Initiation of CNS myelination in the optic nerve is dependent on axon caliber. Cell Rep 25, 544–550.e3. 10.1016/j.celrep.2018.09.052.

24. Bechler, M.E., Byrne, L., and ffrench-Constant, C. (2015). CNS myelin sheath lengths are an intrinsic property of oligodendrocytes. Curr Biol 25, 2411–2416. 10.1016/j.cub.2015.07.056.

25. Bechler, M.E., Swire, M., and ffrench-Constant, C. (2017). Intrinsic and adaptive myelination—A sequential mechanism for smart wiring in the brain. Dev Neurobiol 78, 68–79. 10.1002/dneu.22518.

26. Hernandez, M., Patzig, J., Mayoral, S.R., Costa, K.D., Chan, J.R., and Casaccia, P. (2016). Mechanostimulation promotes nuclear and epigenetic changes in oligodendrocytes. J Neurosci 36, 806–813. 10.1523/JNEUROSCI.2873-15.2016.

27. Jagielska, A., Lowe, A.L., Makhija, E., Wroblewska, L., Guck, J., Franklin, R.J.M., Shivashankar, G. V., and Van Vliet, K.J. (2017). Mechanical strain promotes oligodendrocyte differentiation by global changes of gene expression. Front Cell Neurosci 11. 10.3389/fncel.2017.00093.

28. Segel, M., Neumann, B., Hill, M.F.E., Weber, I.P., Viscomi, C., Zhao, C., Young, A., Agley, C.C., Thompson, A.J., Gonzalez, G.A., et al. (2019). Niche stiffness underlies the ageing of central nervous system progenitor cells. Nature 573, 130–134. 10.1038/s41586-019-1484-9.

29. Urbanski, M.M., Kingsbury, L., Moussouros, D., Kassim, I., Mehjabeen, S., Paknejad, N., and Melendez-Vasquez, C. V. (2016). Myelinating glia differentiation is regulated by extracellular matrix elasticity. Sci Rep 6. 10.1038/srep33751.

30. Ong, W., Marinval, N., Lin, J., Nai, M.H., Chong, Y.S., Pinese, C., Sajikumar, S., Lim, C.T., ffrench-Constant, C., Bechler, M.E., et al. (2020). Biomimicking fiber platform with tunable stiffness to study mechanotransduction reveals stiffness enhances oligodendrocyte differentiation but impedes myelination through YAP-dependent regulation. Small 16. 10.1002/smll.202003656.

31. Shimizu, T., Osanai, Y., Tanaka, K.F., Abe, M., Natsume, R., Sakimura, K., and Ikenaka, K. (2017). YAP functions as a mechanotransducer in oligodendrocyte morphogenesis and maturation. Glia 65, 360–374. 10.1002/glia.23096.

32. Halford, J., Senatore, A.J., Berryman, S., Muñoz, A., Semidey, D., Doan, R.A., Coombs, A.M., Noimany, B., Emberley, K., Emery, B., et al. (2025). TMEM63A, associated with hypomyelinating leukodystrophies, is an evolutionarily conserved regulator of myelination. Proc Natl Acad Sci U S A 122. 10.1073/pnas.2507354122.

33. Wang, Y.Y., Wu, D., Zhan, Y., Li, F., Zang, Y.Y., Teng, X.Y., Zhang, L., Duan, G.F., Wang, H., Xu, R., et al. (2025). Cation channel TMEM63A autonomously facilitates oligodendrocyte differentiation at an early stage. Neurosci Bull 41, 615–632. 10.1007/s12264-024-01338-4.

34. Blaschuk, K.L., Frost, E.E., and ffrench-Constant, C. (2000). The regulation of proliferation and differentiation in oligodendrocyte progenitor cells by αv integrins. Development 127, 1961–1969. 10.1242/dev.127.9.1961.

35. Bechler, M.E. (2019). A neuron-free microfiber assay to assess myelin sheath formation. In Oligodendrocytes: Methods and Protocols Methods in Molecular Biology., D. A. Lyons and L. Kegel, eds. (Springer New York), pp. 97–110. 10.1007/978-1-4939-9072-6_6.

36. Waxman, S.G., and Sims, T.J. (1984). Specificity in central myelination: evidence for local regulation of myelin thickness. Brain Res 292, 179–185. 10.1016/0006-8993(84)90905-3.

37. Almeida, R.G., Czopka, T., ffrench-Constant, C., and Lyons, D.A. (2011). Individual axons regulate the myelinating potential of single oligodendrocytes in vivo. Development 138, 4443–4450. 10.1242/dev.071001.

38. Sharma, K., Schmitt, S., Bergner, C.G., Tyanova, S., Kannaiyan, N., Manrique-Hoyos, N., Kongi, K., Cantuti, L., Hanisch, U.-K., Philips, M.-A., et al. (2015). Cell type– and brain region–resolved mouse brain proteome. Nat Neurosci 18, 1819–1831. 10.1038/nn.4160.

39. Thakurela, S., Garding, A., Jung, R.B., Müller, C., Goebbels, S., White, R., Werner, H.B., and Tiwari, V.K. (2016). The transcriptome of mouse central nervous system myelin. Sci Rep 6, 25828. 10.1038/srep25828.

40. Coste, B., Mathur, J., Schmidt, M., Earley, T.J., Ranade, S., Petrus, M.J., Dubin, A.E., and Patapoutian, A. (2010). Piezo1 and Piezo2 are essential components of distinct mechanically activated cation channels. Science (1979) 330, 55–60. 10.1126/science.1193270.

41. Lappe-Siefke, C., Goebbels, S., Gravel, M., Nicksch, E., Lee, J., Braun, P.E., Griffiths, I.R., and Navel, K.A. (2003). Disruption of Cnp1 uncouples oligodendroglial functions in axonal support and myelination. Nat Genet 33, 366–374. 10.1038/ng1095.

42. Cahalan, S.M., Lukacs, V., Ranade, S.S., Chien, S., Bandell, M., and Patapoutian, A. (2015). Piezo1 links mechanical forces to red blood cell volume. Elife 4. 10.7554/eLife.07370.

43. Acheta, J., Bhatia, U., Haley, J., Hong, J., Rich, K., Close, R., Bechler, M.E., Belin, S., and Poitelon, Y. (2022). Piezo channels contribute to the regulation of myelination in Schwann cells. Glia 70, 2276–2289. 10.1002/glia.24251.

44. Swire, M., Kotelevtsev, Y., Webb, D.J., Lyons, D.A., and ffrench-Constant, C. (2019). Endothelin signalling mediates experience-dependent myelination in the CNS. Elife 8, 1– 23. 10.7554/eLife.49493.

45. Hughes, E.G., Orthmann-Murphy, J.L., Langseth, A.J., and Bergles, D.E. (2018). Myelin remodeling through experience-dependent oligodendrogenesis in the adult somatosensory cortex. Nat Neurosci 21, 696–706. 10.1038/s41593-018-0121-5.

46. Hill, R.A., Li, A.M., and Grutzendler, J. (2018). Lifelong cortical myelin plasticity and age-related degeneration in the live mammalian brain. Nat Neurosci 21, 683–695. 10.1038/s41593-018-0120-6.

47. Call, C.L., and Bergles, D.E. (2021). Cortical neurons exhibit diverse myelination patterns that scale between mouse brain regions and regenerate after demyelination. Nat Commun 12, 4767. 10.1038/s41467-021-25035-2.

48. Hildebrand, C., and Hahn, R. (1978). Relation between myelin sheath thickness and axon size in spinal cord white matter of some vertebrate species. J Neurol Sci 38, 421–434. 10.1016/0022-510X(78)90147-8.

49. Jarjour, A.A., and Sherman, D.L. (2019). Teasing of Ventral Spinal Cord White Matter Fibers for the Analysis of Central Nervous System Nodes of Ranvier. In Oligodendrocytes: Methods and Protocols Methods in Molecular Biology., D. A. Lyons and L. Kegel, eds. (Springer New York), pp. 129–139. 10.1007/978-1-4939-9072-6_8.

50. Snaidero, N., Möbius, W., Czopka, T., Hekking, L.H.P., Mathisen, C., Verkleij, D., Goebbels, S., Edgar, J., Merkler, D., Lyons, D.A., et al. (2014). Myelin membrane wrapping of CNS axons by PI(3,4,5)P3-dependent polarized growth at the inner tongue. Cell 156, 277–290. 10.1016/j.cell.2013.11.044.

51. Chang, K.-J., Redmond, S.A., and Chan, J.R. (2016). Remodeling myelination: implications for mechanisms of neural plasticity. Nat Neurosci 19, 190–197. 10.1038/nn.4200.

52. Etxeberria, A., Hokanson, K.C., Dao, D.Q., Mayoral, S.R., Mei, F., Redmond, S.A., Ullian, E.M., and Chan, J.R. (2016). Dynamic modulation of myelination in response to visual stimuli alters optic nerve conduction velocity. J Neurosci 36, 6937–6948. 10.1523/JNEUROSCI.0908-16.2016.

53. Gibson, E.M., Purger, D., Mount, C.W., Goldstein, A.K., Lin, G.L., Wood, L.S., Inema, I., Miller, S.E., Bieri, G., Zuchero, J.B., et al. (2014). Neuronal activity promotes oligodendrogenesis and adaptive myelination in the mammalian brain. Science (1979) 344. 10.1126/science.1252304.

54. Wake, H., Ortiz, F.C., Woo, D.H., Lee, P.R., Angulo, M.C., and Fields, R.D. (2015). Nonsynaptic junctions on myelinating glia promote preferential myelination of electrically active axons. Nat Commun 6, 7844. 10.1038/ncomms8844.

55. Hines, J.H., Ravanelli, A.M., Schwindt, R., Scott, E.K., and Appel, B. (2015). Neuronal activity biases axon selection for myelination in vivo. Nat Neurosci 18, 683–689. 10.1038/nn.3992.

56. Almeida, R.G., Williamson, J.M., Madden, M.E., Early, J.J., Voas, M.G., Talbot, W.S., Bianco, I.H., and Lyons, D.A. (2021). Myelination induces axonal hotspots of synaptic vesicle fusion that promote sheath growth. Curr Biol 31, 3743–3754.e5. 10.1016/j.cub.2021.06.036.

57. Lewis, A.H., and Grandl, J. (2015). Mechanical sensitivity of Piezo1 ion channels can be tuned by cellular membrane tension. Elife 4. 10.7554/eLife.12088.

58. Lin, Y.C., Guo, Y.R., Miyagi, A., Levring, J., MacKinnon, R., and Scheuring, S. (2019). Force-induced conformational changes in PIEZO1. Nature 573, 230–234. 10.1038/s41586-019-1499-2.

59. Yang, X., Lin, C., Chen, X., Li, S., Li, X., and Xiao, B. (2022). Structure deformation and curvature sensing of PIEZO1 in lipid membranes. Nature 604, 377–383. 10.1038/s41586-022-04574-8.

60. Krasnow, A.M., Ford, M.C., Valdivia, L.E., Wilson, S.W., and Attwell, D. (2018). Regulation of developing myelin sheath elongation by oligodendrocyte calcium transients in vivo. Nat Neurosci 21, 24–30. 10.1038/s41593-017-0031-y.

61. Battefeld, A., Popovic, M.A., de Vries, S.I., and Kole, M.H.P. (2019). High-frequency microdomain Ca2+ transients and waves during early myelin internode remodeling. Cell Rep 26, 182–191.e5. 10.1016/j.celrep.2018.12.039.

62. Auer, F., Vagionitis, S., and Czopka, T. (2018). Evidence for myelin sheath remodeling in the CNS revealed by in vivo imaging. Curr Biol 28, 549–559.e3. 10.1016/j.cub.2018.01.017.

63. Bagheri, H., Friedman, H., Hadwen, A., Jarweh, C., Cooper, E., Oprea, L., Guerrier, C., Khadra, A., Collin, A., Cohen-Adad, J., et al. (2024). Myelin basic protein mRNA levels affect myelin sheath dimensions, architecture, plasticity, and density of resident glial cells. Glia 72, 1893–1914. 10.1002/glia.24589.

64. Iyer, M., Kantarci, H., Cooper, M.H., Ambiel, N., Novak, S.W., Andrade, L.R., Lam, M., Jones, G., Münch, A.E., Yu, X., et al. (2024). Oligodendrocyte calcium signaling promotes actin-dependent myelin sheath extension. Nat Commun 15, 265. 10.1038/s41467-023-44238-3.

65. Collins, H.Y., Doan, R.A., Li, J., Early, J.E., Madden, M.E., Simkins, T., Lyons, D.A., Monk, K.R., and Emery, B. (2025). FBXW7 regulates MYRF levels to control myelin capacity and homeostasis in the adult central nervous system. Nat Commun 16, 7822. 10.1038/s41467-025-62715-9.

66. Pajevic, S., Basser, P.J., and Fields, R.D. (2014). Role of myelin plasticity in oscillations and synchrony of neuronal activity. Neuroscience 276, 135–147. 10.1016/j.neuroscience.2013.11.007.

67. Swire, M., and ffrench-Constant, C. (2019). Oligodendrocyte–neuron myelinating coculture. In Oligodendrocytes: Methods and Protocols Methods in Molecular Biology., D. A. Lyons and L. Kegel, eds. (Springer New York), pp. 111–128. 10.1007/978-1-4939-9072-6_7.

68. Chen, Y., Stevens, B., Chang, J., Milbrandt, J., Barres, B.A., and Hell, J.W. (2008). NS21: Re-defined and modified supplement B27 for neuronal cultures. J Neurosci Methods 171, 239–247. 10.1016/j.jneumeth.2008.03.013.

69. Longair, M.H., Baker, D.A., and Armstrong, J.D. (2011). Simple Neurite Tracer: open source software for reconstruction, visualization and analysis of neuronal processes. Bioinformatics 27, 2453–2454. 10.1093/bioinformatics/btr390.

70. Preibisch, S., Saalfeld, S., and Tomancak, P. (2009). Globally optimal stitching of tiled 3D microscopic image acquisitions. Bioinformatics 25, 1463–1465. 10.1093/bioinformatics/btp184.

