## Supplemental Figures and Text for "CNS Myelin Sheath Lengths Locally Scale to Axon Diameter via Piezo1"

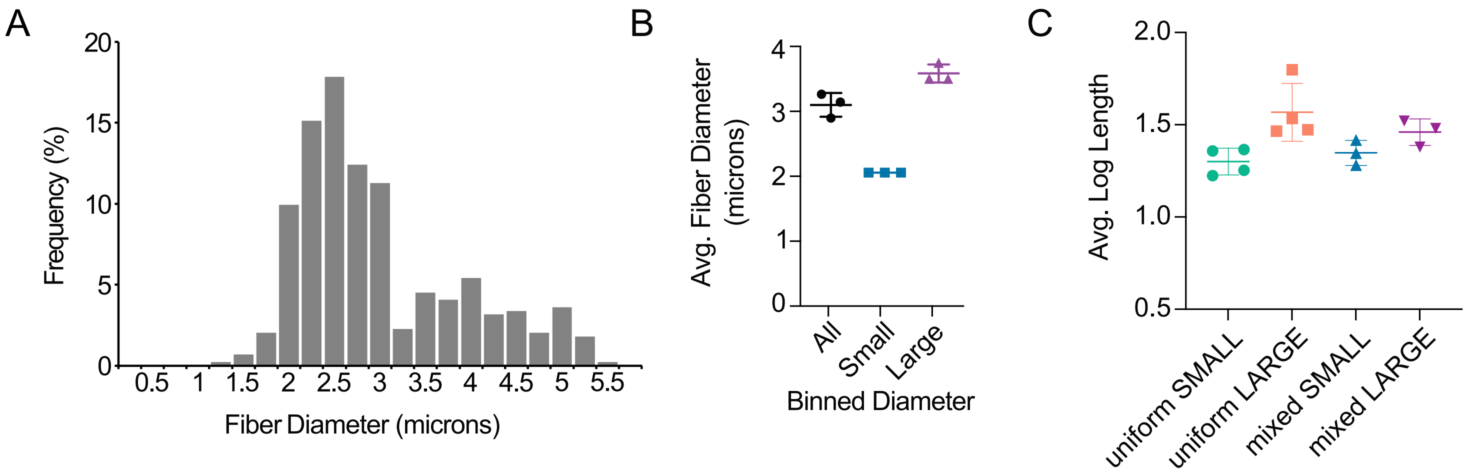
A
Frequency (%)
20
15
10
5
0
0.5
1
1.5
2
2.5
3
3.5
4
4.5
5
5.5
Fiber Diameter (microns)
B
Avg. Fiber Diameter (microns)
4
3
2
1
0
All
Small
Large
Binned Diameter
C
Avg. Log Length
2.0
1.5
1.0
0.5
uniform SMALL
uniform LARGE
mixed SMALL
mixed LARGE

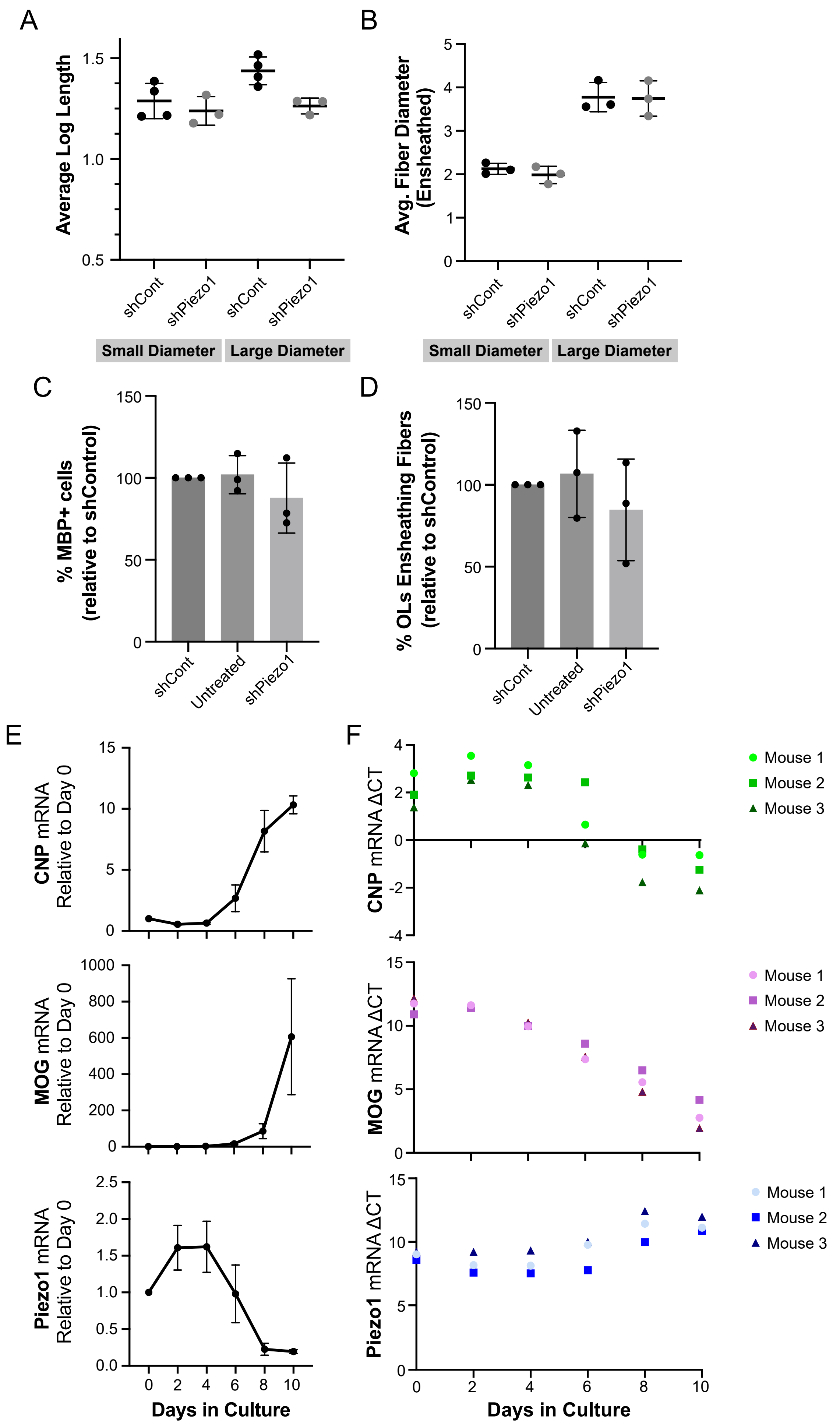

A
Average Log Length
shCont
shPiezo1
shCont
shPiezo1
Small Diameter
Large Diameter
B
Avg. Fiber Diameter (Ensheathed)
shCont
shPiezo1
shCont
shPiezo1
Small Diameter
Large Diameter
C
% MBP+ cells (relative to shControl)
shCont
Untreated
shPiezo1
D
% OLs Ensheathing Fibers (relative to shControl)
shCont
Untreated
shPiezo1
E
CNP mRNA Relative to Day 0
MOG mRNA Relative to Day 0
Piezo1 mRNA Relative to Day 0
Days in Culture
F
CNP mRNA ΔCT
MOG mRNA ΔCT
Piezo1 mRNA ΔCT
Mouse 1
Mouse 2
Mouse 3
Days in Culture

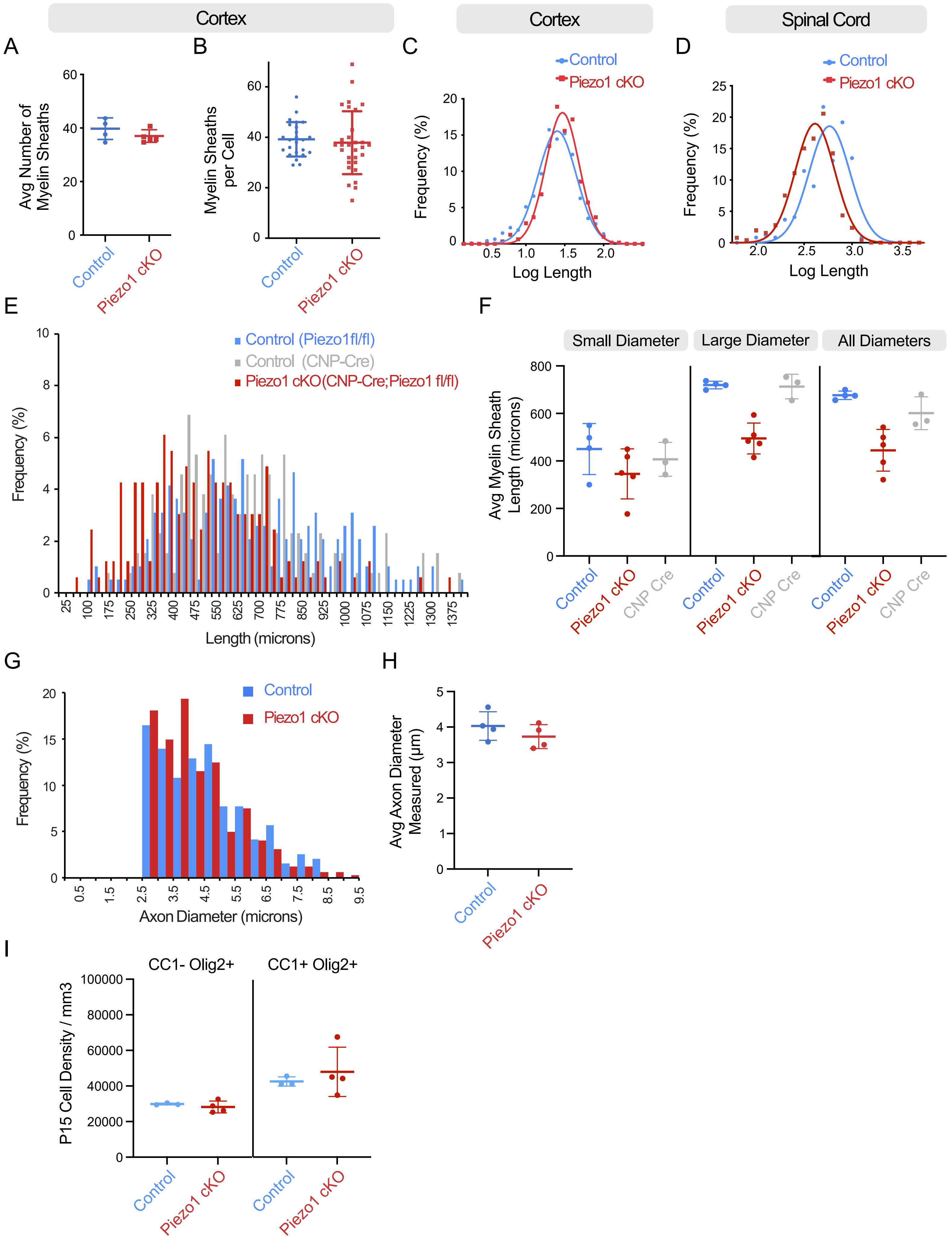

Cortex
Cortex
Spinal Cord
A
B
C
D
Avg Number of Myelin Sheaths
Myelin Sheaths per Cell
Frequency (%)
Log Length
Control
Piezo1 cKO
E
F
Control (Piezo1fl/fl)
Control (CNP-Cre)
Piezo1 cKO(CNP-Cre;Piezo1 fl/fl)
Frequency (%)
Length (microns)
Small Diameter
Large Diameter
All Diameters
Avg Myelin Sheath Length (microns)
CNP Cre
G
H
Axon Diameter (microns)
Avg Axon Diameter Measured (µm)
I
CC1- Olig2+
CC1+ Olig2+
P15 Cell Density / mm3

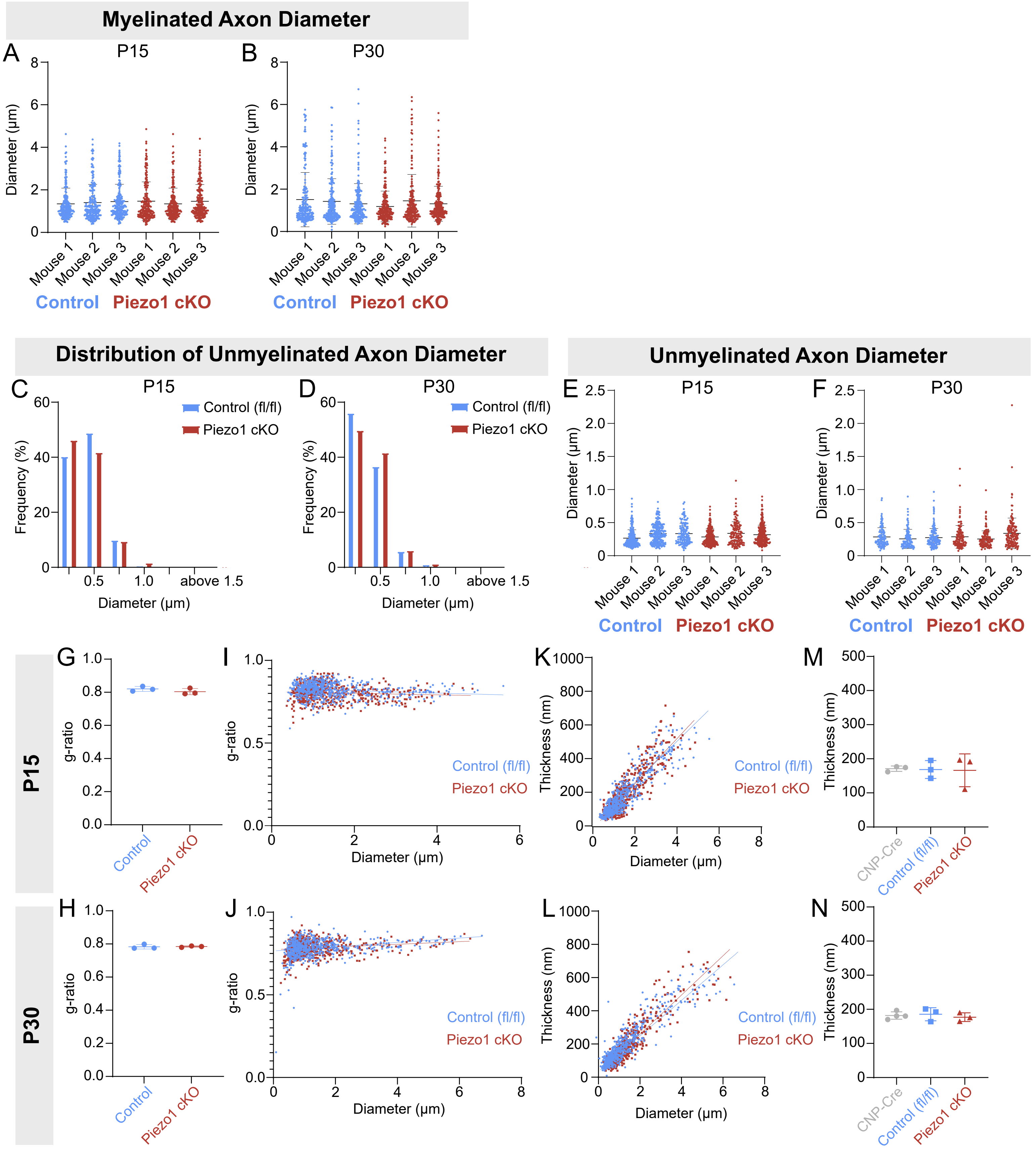

Myelinated Axon Diameter
A
P15
B
P30
Diameter (µm)
Mouse 1
Mouse 2
Mouse 3
Control
Piezo1 cKO
Distribution of Unmyelinated Axon Diameter
C
D
Control (fl/fl)
Piezo1 cKO
Frequency (%)
Diameter (µm)
above 1.5
Unmyelinated Axon Diameter
E
F
G
H
I
J
K
L
M
N
P15
P30
g-ratio
Thickness (nm)
CNP-Cre
Control (fl/fl)

**Supplemental Figure 1. Mixed microfiber diameters and log sheath lengths.**

**A)** Microfiber diameters observed from all mixed diameter cultures. The frequency distribution shows all ensheathed microfiber diameters binned in 0.5-micron increments. **B)** The average microfiber diameter per experiment. All fiber diameters as well as the diameters binned as "small" or "large". **C)** The average log length of myelin sheaths formed in different microfiber cultures. n = 3-4 independent oligodendrocyte microfiber cultures. Error bars = standard deviation.

**Supplemental Figure 2. Piezo1 knockdown in microfiber cultures and expression of Piezo1 mRNA in the mouse oligodendrocyte lineage.**

**A)** The average log length of myelin sheaths formed in the mixed microfiber cultures treated with inducible control shRNA or Piezo1 shRNA, binned into myelin sheaths formed on small diameter (<2.5 micron) or large diameter (> 2.5 micron) fibers. **B)** Microfiber diameters observed in knockdown culture experiments. The average microfiber diameter per experiment. Diameters binned as "small" or "large". n = 3-4 independent oligodendrocyte microfiber cultures. **C)** OL differentiation (% MBP+ cells) relative to control shRNA treated conditions in knockdown microfiber experiments. **D)** Proportion of MBP+ OLs wrapping microfibers relative to control shRNA-treated conditions. n = 3 independent oligodendrocyte cultures (different pooled rat litters) on microfibers. **E)** Quantitative PCR of mouse oligodendrocyte lineage cells acutely isolated (Day 0) through differentiation in culture (up to Day 10 in culture). mRNA expression levels are shown relative to Day 0 (at time of PDGFRa+ mouse cell isolation) after normalization to housekeeping gene GAPDH. CNP and MOG levels rise dramatically and Piezo1 mRNA levels decrease after 8 days in culture. **F)** Delta Ct values (amplification cycle relative to GAPDH) across time points for three mice. CNP levels are already robustly present but increase further when MOG levels rapidly rise. Piezo1 mRNA levels are relatively low and decrease further in the last two days in culture. n = cells isolations from 3 different mice. Error bars =

standard deviation.

**Supplemental Figure 3. Piezo1 is important for the elongation of myelin sheaths on large diameter axons in vivo.**

**A)** The average number of myelin sheaths formed by single OLs in the mouse frontal cortex, layers II-III at P30. p = 0.28, Welch's corrected t-test, n = 4 - 5 mice per genotype. **B)** Number of myelin sheaths formed by single OLs from all the mice. **C)** Log gaussian curve showing the distribution of myelin sheath lengths (log length) in the mouse frontal cortex at P30. p = 0.56 Welch's corrected t-test of mean log lengths (n = 4 - 5 mice). **D)** Log gaussian curve showing the distribution of myelin sheath lengths (log length) on teased spinal cord axons at P30, with a shift in sheath length distributions. **E)** The distribution of myelin sheath lengths in the mouse spinal cord from Piezo1 CKO mice, alongside both control groups: floxed control littermates and CNP-Cre control mice. Distribution of all myelin sheath lengths. **F)** Mean sheath lengths per mouse at P30 on all axon diameters as well as those binned by axon diameters less than or greater than 2.5 microns. Piezo1 cKO mice compared against both floxed littermate controls (Control) or CNP-Cre controls, p = 0.002 for all axons, p = 0.33 for small diameter, p = 0.0001 for large diameter, one-way ANOVA. n = 3 - 5 mice per genotype, at least 62 myelin sheaths measured per mouse. **G)** Distribution of the large diameter axons with measured myelin sheath lengths from teased spinal cords from 4 - 5 mice measured in D-F. **H)** Mean axon diameter per mouse. Measurements taken from all spinal cord axons measured in Figure 3. p = 0.30, Welch's corrected two-tailed t-test. **I)** Density of all immature (CC1-) and mature (CC1+) oligodendrocyte lineage cells at P15 in the mouse spinal cord. n = 3 - 4 mice per genotype, p = 0.39 and p = 0.49, Welch's corrected t-test. Error bars = standard deviation.

**Supplemental Figure 4. Myelin ultrastructural data: calculated g-ratio, individual**

**thickness measurements and diameter distributions from myelinated and unmyelinated axons.**

**A, B)** Distribution of myelinated axon diameter by mouse at P15 and P30. **C, D)** Frequency distribution of unmyelinated axon diameters at P15 and P30. **E, F)** Distribution of unmyelinated axon diameter by mouse at P15 and P30. **G-J)** Equivalent g-ratios for thickness data. g-ratios were calculated from the axon perimeter measurements and the thickness of the myelin sheath (g-ratio = axon diameter / (axon diameter + (thickness x 2))). **G, H)** Average g-ratio at P15 (t-test with Welch's correction, p = 0.31) and P30 (test-test with Welch's correction, p = 0.89). **I, J)** Scatter plot of calculated g-ratio at P15 (simple linear regression, $R^2$ = 0.01) and P30 (simple linear regression, $R^2$ = 0.28). **K, L)** Scatter plot of myelin thickness measurements at P15 (simple linear regression, $R^2$ = 0.79) and P30 (simple linear regression, $R^2$ = 0.84). **M,N)** Average sheath thickness including CNP-Cre controls at P15 (cKO vs CNP-Cre, p = 0.87) and P30 (p = 0.62). p values from t-tests with Welch's correction between two genotypes.
